# Environmental stressors induced strong small-scale phenotypic differentiation in a wide-dispersing marine snail

**DOI:** 10.1101/2020.10.06.327767

**Authors:** Nicolás Bonel, Jean-Pierre Pointier, Pilar Alda

## Abstract

Heterogeneous environments pose a particular challenge for organisms because the same phenotype is unlikely to perform best regardless of the variety of encountered stressors. To understand how species meet this challenge, we investigated the extent to which contrasting environmental pressures induced ecological and phenotypic responses in a natural population of a wide-dispersing marine snail at a small spatial scale. We analyzed several traits of *Heleobia australis* (Rissooidea: Cochliopidae) collected from heterogeneous, but highly connected, habitats from the intertidal area of the Bahía Blanca estuary, Argentina. We also conducted molecular analyses by amplifying the COI gene in individuals sampled from each habitat. We found that sympatric subpopulations of *H. australis* exhibited a strong phenotypic divergence in shell characters and body weight in response to thermal, saline, and dehydration stress, crab predation risk, and parasitic castrators. We proved that this differentiation occurred even early in life as most of the characters observed in juveniles mirrored those found in adults. We also found a divergence in penis size in snails collected from each habitat and raised in common garden laboratory conditions. The molecular analyses confirmed that the individuals studied constituted a single species despite the strong phenotypic differences among subpopulations. The small-scale phenotypic differentiation suggests that *H*. *australis* experienced a fine-grained environment where conditions imposed by different sources of stress favored the expression of beneficial traits. We discuss the role of plasticity in shaping adaptive phenotypic responses that increase the likelihood of persistence of subpopulations facing environmental stress conditions.

## 1. INTRODUCTION

If organisms have difficulties in adapting to human-induced global change and fail to track projected environmental changes, populations become vulnerable to decline and extinction (Hill et al. 2011, Hoffmann & Sgrò 2011). Phenotypic plasticity, a common feature in nature across many taxa, is the ability of a single genotype to express different phenotypes in dissimilar biotic or abiotic environments (Travis 1994, West-Eberhard 2003). Many studies indicate the role of adaptive plasticity in shaping phenotypic responses as an effective mechanism that can greatly improve the conditions for persistence of populations facing abrupt environmental shifts by facilitating rapid adaptation to the new selection pressures (Price et al. 2003, Ghalambor et al. 2007, Chevin & Lande 2009). Adaptive plasticity places populations close enough to the new favored phenotypic optimum for directional selection to act. This provides the first step in adaptation to new environments on ecological time scales as long as costs to maintaining and/or producing a plastic response are not substantially large, thus constraining the evolution of plasticity (Price et al. 2003, Ghalambor et al. 2007, Chevin & Lande 2009). In this sense, adaptive phenotypic plasticity is crucial for the long-term persistence of populations facing new environmental stress brought about by human-induced global change (Hill et al. 2011).

Temperate marine habitats, in particular the intertidal zone, exhibit great variability in environmental factors mainly driven by the fall and rise of the tides, creating areas on the shore that are alternately immersed and exposed (Denny & Paine 1998, Bourdeau et al. 2015). The transitional nature of the intertidal habitat from marine to terrestrial conditions strongly influences the physiology and the ecology of intertidal organisms due to, for instance, the increase in environmental harshness (e.g. desiccation, extremes in temperature and salinity) along the vertical zonation of the intertidal area (Vermeij 1972, Denny & Paine 1998, Mouritsen et al. 2018). From this perspective, intertidal organisms, such as aquatic gastropods, are likely to exhibit increased phenotypic plasticity in response to such environmental selective pressures (Berrigan & Scheiner 2004).

Phenotypic variation in shell traits is particularly widespread in gastropods (Bourdeau et al. 2015) and their shells offer an easily measured and permanent record of how the organism responded to local abiotic (e.g. water chemistry, temperature) and biotic (e.g. predation, parasitism) agents (Dillon et al. 2013). Major environmental factors such as prolonged submersion times, desiccation, high temperature, and extreme salinity across the vertical zonation create contrasting pressures that give rise to opposite shell trait responses (Struhsaker 1968, Johannesson & Johannesson 1996). For instance, prolonged submersion times enhance foraging times and the absorption of calcium carbonate from water leads to higher shell growth rates, resulting in larger and narrower shells and increased body mass <u>(Vermeij 1973,</u> review by <u>Chapman 1995)</u>. By contrast, snails from upper intertidal habitats that are periodically exposed to hot and dry conditions and have only a limited time each day to feed, they exhibit the opposite shell and body mass responses. Their characteristic smaller aperture size has been correlated to an increase in resistance to desiccation in upper intertidal habitats (Vermeij 1973, Chapman 1995, Melatunan et al. 2013).

Predators are a significant contributor to heterogeneity across the intertidal zone. They are often distributed patchily in time and space, creating a steep selection gradient in predation risk, which can also promote adaptive shell responses (Appleton & Palmer 1988, Palmer 1990, Boulding & Hay 1993, Hollander et al. 2006, Bourdeau 2011). The adaptive value of predator-induced plasticity primarily linked to shell defenses are not limited to shell thickening (more resistant to crushing), as snails are capable of plastically altering different aspects of their shell shape (i.e. increased globosity; lower shell length to width ratio) or aperture (i.e. more elongated; higher aperture length to width ratio). These adaptations prevent the crab from pulling the soft parts out of the shell (Johannesson 2003, Dillon et al. 2013, Bourdeau et al. 2015). However, not all phenotypic plasticity triggered by environmental conditions is adaptive (Ghalambor et al. 2007). Some plastic trait responses, usually those imposed by the biochemistry and physiology of the organism, can be reversed over short time scales. This is not the case for developmental plasticity, which tends to be irreversible or takes longer to be reversed (Pigliucci et al. 2006). Parasites, for instance, can induce morphological, behavioral, and physiological change of individual snail hosts, and thereby influence several aspects of host life history that can significantly alter size structure, demography, resource use, and intra and interspecific interactions of the host population (Fredensborg et al. 2005, Miura et al. 2006, Mouritsen et al. 2018). They have been also reported to induce changes in microhabitat choice (Curtis 1987) and body size in snails (Mouritsen & Jensen 1994, Probst & Kube 1999, Levri et al. 2005, Miura et al. 2006, Alda et al. 2010). Such pressure exerted by parasites also differs along the intertidal zone (Smith 2001, Alda et al. 2010, 2019) and can cause important shifts in the expression of host phenotypic traits, creating pronounced phenotypic differences between infected and uninfected hosts (Poulin & Thomas 1999, Fredensborg et al. 2006).

Despite the fact that the intertidal area spans a strong selective gradient, small-scale phenotypic differentiation and adaptation of wide-dispersing organisms has been understudied (review by Sanford & Kelly 2011). These contrasting conditions along the vertical distribution pose a particular challenge for wide-dispersing intertidal organisms: the same phenotype is unlikely to perform best across the variety of encountered stressors. As a result, some particular strategies may emerge: (i) a generalist phenotype (probably with maladaptation), (ii) phenotypic plasticity promoting different phenotypes under different conditions, or (iii) a local genetic differentiation, contributing to the phenotypic variation between habitats, favored by restricted gene flow through habitat selection (and probably partial genetic isolation leading to ecotype formation). For instance, limited gene flow can occur when habitat conditions disrupt larval dispersal thereby increasing localized recruitment (Struhsaker 1968, Johnson et al. 1994, Parsons 1997, Levin et al. 2006) where individuals would be subjected to local biotic and abiotic stressors (Johannesson & Johannesson 1996, Westram et al. 2021). This can in turn lead to ‘phenotype-environment mismatch’ (reviewed in Marshall et al. 2010). To understand how species meet this challenge, the first step is to observe phenotypic variation and its association in the field with microhabitats characterized by different sources of stress. The next step is to explore whether such phenotypic variation results from a direct impact of stress (e.g. trait shift induced by parasites) or from ecotypes that find alternative niches (e.g. different microhabitats), thus favoring reproductive isolation between ecotypes (Westram et al. 2018).

Here we tested how contrasting biotic and abiotic conditions across the intertidal gradient affected density and induced different phenotypic responses at a small spatial scale among individuals from subpopulations of a wide-dispersing intertidal snail. We expected a decreased snail density and a decrease in trait mean values in snails inhabiting areas with high physical stress conditions (thermal, saline, and dehydration stress), crab predation risk, or parasite pressure. To test this, we collected, weighed, dissected, and measured shell traits for individuals of the mud snail *Heleobia australis* (d’Orbigny, 1835) (Rissooidea: Cochliopidae) over all four seasons in a year from three distinct, but highly connected, habitats from the intertidal area of the Bahía Blanca estuary, Southwestern Atlantic, Argentina. We then collected juveniles from each habitat and raised them in common garden laboratory conditions until they were adults to investigate if genital morphology varied. Finally, we explored whether individuals living in these contrasting habitats belong to the same species, and not to a species complex, by amplifying the cytochrome oxidase subunit 1 (COI) gene. We found clear evidence of small-scale phenotypic differentiation in all traits, likely as a result of conditions imposed by strong biotic and abiotic stressors across the vertical distribution of the intertidal area.

## 2. MATERIAL AND METHODS

### 2.1. Study species

The intertidal mud snail *Heleobia australis* (previously known as *Littoridina australis*) has a wide geographic range inhabiting marine and estuarine ecosystems from tropical to temperate regions (from Brazil: 22° 54’ S, to Argentina: 40° 84’ S; De Francesco & Isla 2003) and is the most common benthic macrofaunal species in the estuaries and coastal lagoons (De Francesco & Isla 2004). In the Bahía Blanca estuary, there are no organisms that compete with *H*. *australis* as this snail species is the dominant gastropod (Elías et al. 2004). It is a gonochoristic species with internal fertilization (Neves et al. 2010). Mature females lay egg capsules, containing one fertilized egg (Neves et al. 2010), preferably on shells of conspecific individuals, but they may also be laid on shells of other species, on sand grains, or algae. These eggs later develop into a veliger larva that exhibits a short pelagic larval life (Neves et al. 2010), on average 10±3 days in standard laboratory conditions (12:12 photoperiod, 25 °C, water salinity 30 PSU, and *ad libitum* food in the form of boiled ground lettuce; Bonel, *unpublished data*). Adult snails attain a maximum shell size ranging from 7 to 8 mm with an average shell length of 5.2 mm (0.5 SD; this study), a lifespan of 2.9 years, and two well-defined recruitment episodes in a temperate climate (De Francesco & Isla 2004, Carcedo & Fiori 2012). However, in tropical and subtropical areas breeding and recruitment occur year-round (Neves et al. 2010) suggesting that it is likely that *H*. *australis* have shorter generation times. This species is capable of creating an air bubble inside its pallial cavity, allowing juveniles and adults to temporarily float in the water column and be carried along by the tide and wind driven current (Echeverría et al. 2010), increasing its dispersal potential. *H*. *australis* is also an obligate intermediate host of a large diversity of parasites that infect several other host species. One study reported that *H*. *australis* hosted 18 larval trematode species, 16 of which were from the Bahía Blanca estuary (Argentina), and all but one of which were parasitic castrators. The mechanism of these castrators is an individual worm that fills up the digestive gland and gonad of the snail host (Alda & Martorelli 2014).

### 2.2. Study area and (a)biotically heterogeneous habitats

This study was conducted in the Villa del Mar saltmarsh-mudflat located in the middle reaches of the Bahía Blanca estuary, Argentina (38° 51’ S – 62° 07’ W). The intertidal mudflat extends for more than 1 km across the tidal gradient. Its topography is characterized by a gently sloping ramp on the seaward side (Pratolongo et al. 2009) and it is affected by strong semidiurnal tides and high seasonal variation (Perillo et al. 2001). Close to the mean high tide level, marshes covered by cordgrass (*Spartina alterniflora*) form a narrow, 150 m wide strip of vegetation followed by a mudflat area with no vegetation (Pratolongo et al. 2010). During low tide, these upper areas of the intertidal zone are subjected to high desiccation and temperature and salinity fluctuations, as the tide takes *ca*. 10 hours to cover them (P. Pratolongo, *personal communication*; 26 February 2019). Low areas located close the seaward edge remain covered by water during low tide.

The existence of a tidal cycle exposes *H. australis* to extreme, but predictable, changes in abiotic conditions (at least twice every day), likely determining their vertical distribution (zonation patterns). Snails in the upper intertidal zone must remain quiescent for long periods of time during daylight hours or during low tide, as they are exposed to aerial conditions for longer periods than individuals in the lower intertidal zone that remain cover by water during low tide. The intertidal zone can therefore be characterized in three distinct, but spatially adjacent, habitats: flats, marshes, and pans. Flats and marshes are located in the upper zone and they drain at low tide, though flats are free of vegetation and marshes are covered by cordgrass. Pans are free of vegetation but remain covered by water during low tide and are located close the seaward edge. Thermal, saline, and dehydration stress are strong selective forces occurring mainly in the upper intertidal area (flats and marshes; Denny & Paine 1998, Bourdeau et al. 2015) whereas these inducing agents are weaker in pans, which exhibit low environmental stress condition (Fig. 1). The ecology and selective pressures at each habitat are different. However, the lack of geographical separation, the effect of strong semidiurnal tides, and the species’ high dispersion potential facilitate physical encountering among individuals within the intertidal area. We can therefore consider that subpopulations of *H*. *australis* are sympatrically distributed (review by Futuyma & Mayer 1980, but see Butlin et al. 2008).

**Figure 1.**
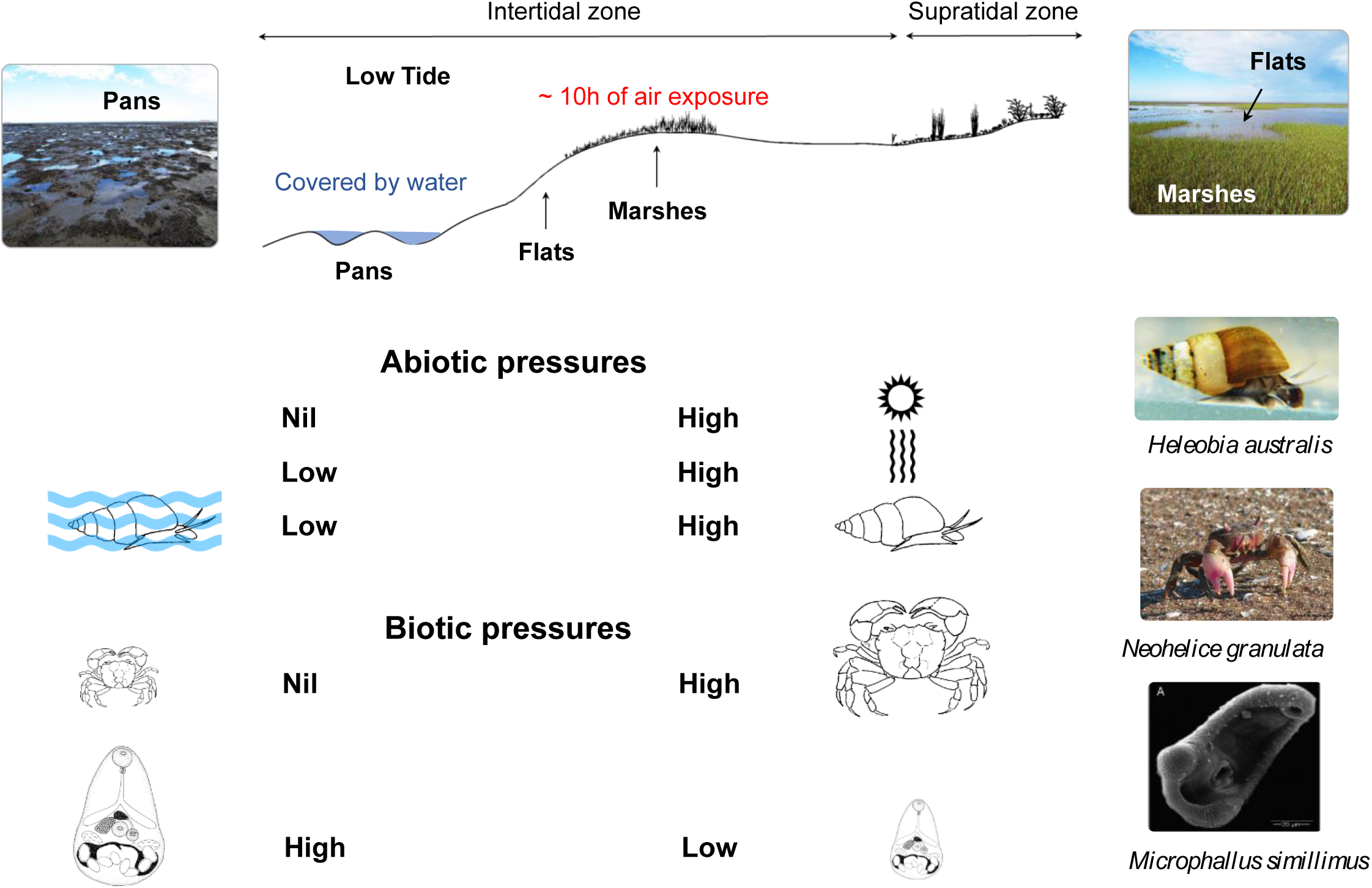
Contrasting habitat conditions at the intertidal zone of the Bahía Blanca estuary (Villa del Mar), Argentina. Three distinct habitats can be characterized: flats, marshes, and pans. Flats and marshes are located in the upper zone and they drain at low tide, though flats are free of vegetation and marshes covered by cordgrass (*Spartina alterniflora*). Pans are free of vegetation but remain covered by water during low tide and are located close the seaward edge. Thermal, saline, and dehydration stress are strong selective forces occurring mainly in the upper intertidal area (flats and marshes) whereas these inducing agents are weaker in pans, which exhibit low environmental stress condition. Biotic agents also vary along the vertical distribution of the intertidal zone. Predation risk is higher in the upper area (marshes) mainly driven by the grapsid burrowing crab *Neohelice granulata*, which is commonly found in high abundances in marshes, though it uses the entire intertidal zone. Parasite pressure also differs along the intertidal zone. The prevalence (percentage of individuals infected) of trematodes is higher in the lower area of the intertidal zone (pans), where parasite infection is predominately caused by one extremely prevalent trematode, *Microphallus simillimus*.

Biotic agents also vary along the vertical distribution of the intertidal zone (Fig. 1). On the one hand, the grapsid crab *Neohelice granulata* is one of the most abundant macroinvertebrates of intertidal areas of the SW Atlantic estuaries where it commonly inhabits the upper vegetated area (marshes) but uses the entire intertidal zone (Spivak et al. 1994, Alvarez et al. 2013, Angeletti et al. 2014, Angeletti & Cervellini 2015). This burrowing crab could exert a strong pressure on *Heleobia australis* since it can drastically reduce snail density in vegetated areas due to an intense bioturbation activity (Spivak et al. 1994, Alvarez et al. 2013, Angeletti et al. 2014, Angeletti & Cervellini 2015) and/or through snail predation (D’Incao et al. 1990, Barutot et al. 2011). Although the main predator in the intertidal mudflat are crabs, there are other potential predators such as snail-eating birds (e.g. *Charadrius* spp.) that occasionally feed on *H. australis* (Martorelli 1991, Ieno et al. 2004, Meerhoff et al. 2013). For instance, in the study area, *H. australis* has been reported to be one of the 39 dietary items consumed by the White-Backed Stilt (Petracci & Delhey 2005). On the other hand, parasite pressure also differs along the intertidal zone. Trematode infection not only inevitably leads to snail castration, reducing its fitness to zero (Alda et al. 2019), but also gives rise to smaller shell-sized morphs for infected snails (Alda et al. 2010). The prevalence (percentage of individuals infected) of trematodes is higher in the lower area of the intertidal zone, where parasite infection is predominately caused by one extremely prevalent trematode, *Microphallus simillimus.* Such pressure is stronger in pans than in flats and marshes because prolonged submersion times allow snails to increase time spent foraging and thus ingesting parasite eggs (Alda et al. 2019).

### 2.3. Field sampling and laboratory procedure

We sampled individuals of the intertidal mud snail *Heleobia australis* in summer (March 6), autumn (July 15), winter (September 20), and spring (December 7) of 2012 from the study area. We performed intense sampling in three contrasting intertidal habitats from the Villa del Mar saltmarsh-mudflat because our aim was to test for phenotypic variation in *H. australis* at a small-spatial scale (< 1 km) and because marine environmental conditions in this location are representative of conditions found at other sites from the Bahía Blanca estuary (Piccolo & Hoffmeyer 2004). We followed a simple random sampling design by area for polygon study areas. We used QGIS 1.7 (https://qgis.org) to define the sampling zone and to randomly distribute the points where the circular quadrats were placed. Specifically, during low tide and from each biological zone from the intertidal area (flats, marshes, and pans), we sampled snails by laying down nine circular quadrats with 10 cm diameter and 2 cm deep (Area = 78.5 cm^2^) to get a representative sample of each zone (e.g. Milroy 2015). Snails were sieved and washed from the sediment through a 1-mm mesh, and immediately transported alive to the laboratory in 0.5-l plastic bottles filled with *in situ* water where they were kept in aquaria and fed *ad libitum* with flake fish food. We repeated this procedure for each sampling date.

Once in the lab, we counted the total number of snails in each of the nine samples (i.e. samples A to I) taken from each habitat (flats, marshes, or pans) and sampling date. From each sample, we arbitrarily decided to subsample 60% of individuals from each sample to check for parasite infection, 30% for shell and body weight, and 10% for parasite biomass. Because *Heleobia australis* is rather a small snail (mean shell length = 5-6 mm), it is difficult to visually assess whether individuals are phenotypically different from one another. Thus, as the probability of choosing each individual was equal, we assumed that subsampling was done at random. Overall, our large dataset consisted of 6,250 individuals, which were used to account for infected and uninfected snails (hereafter infection status), and the remaining uncrushed snails were used to estimate snail shell and body weight (n = 3,057; see below) and parasite biomass (n = 1,060; data not showed in this study). All 10,367 snails were used to estimate snail density and were also photographed using a camera attached to a dissecting microscope to analyze shell and aperture morphometrics (see below).

### 2.4. Snail density and infection status

To test how contrasting habitat conditions affected snail density, we counted all sampled individuals per habitat and sampling date to analyze spatial and temporal variation of snail density. Then, to determine infection status, snails from each habitat were crushed using a mortar and a pestle, tissue was examined under a dissecting microscope, and trematodes were identified under a compound microscope following Alda & Martorelli (2014). Previous studies show that the most prevalent parasite in the study area is *Microphallus simillimus* (Microphallidae), which makes up 87% of the overall prevalence (Alda & Martorelli 2014, Alda et al. 2019). This parasite has an abbreviated life cycle (the same snail host can serve as the first- and second-intermediate host), meaning that once snails ingest parasite eggs, the metacercariae encyst within the sporocyst in the infected snail, thereby maximizing transmission success (Alda & Martorelli 2014). One possible approach to analyze the contribution of parasites to the phenotypic variance in host populations is to compare phenotypic responses of uninfected and infected hosts maintained under identical conditions (Poulin & Thomas 1999). To test for this, we thus only considered snails infected by the most prevalent trematode *M. simillimus* and also from pans (the habitat with the highest prevalence; Alda et al. 2019), which implies that both infected and uninfected snails grew in the same environmental conditions. By doing so, we can ensure that differences in phenotypic responses can only be attributed to *M. simillimus* effect and not caused by other trematodes or habitat stress conditions.

As older hosts have greater cumulative risk of infection than do young hosts, each sample should be standardized by age and collected at the same time from an area in which hosts are likely to mingle and where they have experienced relatively uniform risks of infection (Lafferty et al. 1994). We therefore identified age cohorts by means of the length-frequency distributions in each sampling date and habitat and removed those individuals outside the lower 95% confidence limit of the cohort with larger individuals (see Text S1 in the appendix for details; Fig. S1).

### 2.5. Shell and aperture morphometric

To analyze variation in shell morphometric across habitat conditions and infection status, we measured four linear variables from each photograph using ImageJ software: shell length (SL), shell width (SW), aperture length (AL), and aperture width (AW).

We chose the linear morphometrics approach over the geometric morphometrics analysis because the landmark-based approach is more time-consuming and labor-intensive, thus reducing the number of individuals than can be processed (n < 600; e.g. Carvajal-Rodríguez et al. 2005, Conde-Padín et al. 2007, Guerra-Varela et al. 2009, Dillon et al. 2013, Larsson et al. 2020). However, the approach followed in our study allowed us to measure more than 10,000 snails. Similar to some other caenogastropod species (e.g. *Littoridina* sp.), the shell of *H. australis* is generally small and conical, with the axis of coiling lying at an angle of about 45° above the plane of the elliptical aperture (Vermeij 1973, Hershler & Thompson 1992). We estimated the shell volume as a proxy of shell size by calculating the volume of a cone (or double cone) because it is a more reliable measure to describe shell size than only considering a linear measurement such as shell length:

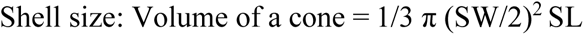

To analyze the shell shape, we estimated the SL to SW ratio. Likewise, we analyzed aperture shape by calculating the AL to AW ratio. We used AL and AW to calculate aperture size by calculating the area of an ellipse:

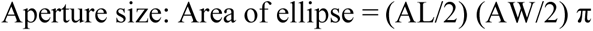

### 2.6. Shell thickness and body mass

To test whether snails subjected to a stronger predation pressure exhibit thicker shells, (considered to be a shell defense trait against predators as it is more resistant to crushing; Johannesson 2003, Dillon et al. 2013, Bourdeau et al. 2015) and whether contrasting environmental stress conditions were affected body mass, we estimated and compared shell weight and ash-free dry-weight among habitats. We used a subsample of individuals (n = 70) representative of the population shell length-frequency distributions that were obtained previously for all individuals sampled on each date and habitat. We removed sediment and epibiota with a scouring pad before weighing and measuring the organic content of the snail as ash-free dry weight (AFDW, calculated as the difference between the dry weight and the weight of the incombustible component of the shell), which was considered as body mass. To achieve this, we dried snails individually in porcelain crucibles for 48 h at 60 °C, weighed them with a digital scale (precision 0.1 mg), ashed the snails for 5 h in a muffle furnace at 500 °C and then reweighed them (Bonel & Lorda 2015). The ash shell weight (the incombustible component of the shell) was considered as proxy of shell thickness (e.g. Palmer 1981) and used to estimate and compare its variation among individuals along the vertical gradient of the intertidal area.

### 2.7. Variation in penis size

To test for differences in genital morphology, we analyzed variation in the penis size and morphology of individuals from contrasting habitats conditions but raised in common garden laboratory conditions. Genital morphology is one of the most widely-used trait when studying the internal anatomy of gastropods, especially among cochliopids (Liu et al. 2001). Reproductive anatomy is very useful to delineate species since they tend to show more variation among than within species and penis morphology, in particular, is used when delineating *Heleobia* species (Gaillard & Castellanos 1976). We sampled snails of *H*. *australis* from the three distinct habitats of the intertidal area of the Villa del Mar saltmarsh-mudflat in November (Spring) of 2017. Snails were transported to the lab and they were split into adults and juveniles based on their shell-length distribution (as mentioned above). Adults were preserved in 96% alcohol and used to conduct molecular analysis. Because field-collected snails may retain the influence of environmental conditions experienced prior to collection even in early life stages (Sanford & Kelly 2011), we tried to minimize this potential effect by considering only the smallest individuals sampled from each habitat (mean shell length, SD: 4.15 ± 0.36 mm). Juveniles were placed in three different aquaria per habitat to prevent tank effects. The snail abundance at the onset of the experiment was similar in each set of aquaria to minimize potential density-dependent effects (flats: n = 38.3 ± 0.6; marshes: n = 43.7 ± 0.6; pans: n = 46.7 ± 0.6). Juveniles were kept in common garden laboratory conditions (12:12 photoperiod, 25 °C, water salinity 30 PSU, and *ad libitum* food in the form of boiled ground lettuce) for 21 months, which ensured that all individuals considered in the analysis were adults. Prior to dissection, we collected, on average, 30 snails from each aquaria and preserved them in the Railliet–Henry’s solution (930 ml distilled water, 6 g NaCl [0.85%], 50 ml formaldehyde [37%], 20 ml glacial acetic acid). Then, they were all dissected under the dissecting microscope and the reproductive system of *ca*. 15 males per aquarium was drawn using a camera lucida attachment (Pointier et al. 2004). These were scanned and the penis surface area was calculated using the ImageJ software. Shells were photographed and analyzed as mentioned above to estimate shell size and shape.

### 2.8. COI analysis

To confirm that individuals living in these contrasting habitats belong to the same species and not to a species complex, we conducted molecular analyses by amplifying the cytochrome oxidase subunit 1 (COI) gene in 10 individuals sampled from each of the three sites. Then, we built a gene tree and a haplotype network to verify if individuals gather in clusters depending on the habitat they come from. We also applied a species-delimitation method: Automatic Barcode Gap Detection (ABGD, Puillandre et al. 2012). ABGD detects barcode gaps, which can be observed whenever the divergence among organisms belonging to the same species is smaller than divergence among organisms from different species (see Text S2 in the appendix for specific details on these procedures).

### 2.9. Statistical analyses

To test for spatial and temporal variation of snail density, we performed a two-way ANOVA including habitats (flats, marshes, and pans) and seasons (summer, autumn, winter, spring) as fixed effects, and their interaction. We transformed density (natural-log) data to meet assumptions of normality and homoscedasticity.

To test for phenotypic variation in shell characters, we defined three categories (cat.) of snails that comprised: (*i*) uninfected adults, (*ii*) infected adults, and (*iii*) uninfected juveniles. These three categories (cat. *i*–*iii*) were considered to test whether contrasting (a)biotic stress conditions (environmental stress and crab predation) induced variation in shell traits (morphometric and thickness) and body mass. To evaluate the effect exerted by the trematode *Microphallus simillimus* on shell morphometry of *Heleobia australis* from pans (the habitat with the highest prevalence), we created a fourth category that included uninfected and infected adult snails (cat. *iv*). We performed four Principal Components Analyses (PCAs) to visualize the main components of the morphological shell variation for each snail category. Then, we tested for differences in shell morphology (size and shape of the shell and aperture) by fitting independent linear mixed models (LMMs), with a Gaussian error distribution, for the above-mentioned four categories (cat. *i*–*iv*).

To test the effect of habitat conditions on shell traits (cat. *i*–*iii*), we considered habitat as a fixed effect. To test for morphometric differences between infected and uninfected individuals (cat. *iv*), we included infection status (infected and uninfected) as a fixed effect as we only considered individuals from one habitat (i.e. same habitat conditions for infected and uninfected snails). For all the four categories (cat. *i*–*iv*), we included sex (male and female) as a fixed factor (and its interaction with habitat or infection status). We added sampling season as a random factor in all models as it can explain a significant portion of the variance in measured variables. Likewise, we also considered shell size as a covariate when testing for shell and aperture shape, but the models were all run with and without this effect to check the extent to which the effects were mediated by shell size. We also incorporated shell size as a covariate when testing for shell and body weight (by means of LMMs) and estimates were statistically corrected for variation that could be explained by the covariate. Statistical significance of the fixed effects was obtained from model comparisons using likelihood-ratio tests. Random effects were separately tested using chi-square likelihood-ratio tests with the corrections indicated by Zuur (2009). To test for differences in penis size (area of the penis), we performed a linear model with habitat as a factor and we added shell size (volume) as a covariate, which was used to statistically correct estimates for variation explained by the covariate. All individuals analyzed were free of parasites, meaning that the differences found are not due to parasite effect.

Post-hoc tests were performed when the effects were significant (*P* <0.05), using Holm-Bonferroni correction for multiple testing to compare their effects. Values are given as means ± 1 SE unless otherwise stated. All analyses and figures were performed with R v.3.3.3 packages *lme4* (Bates et al. 2014), *nlme* (Pinheiro et al. 2017), *car* (Fox et al. 2011), *effects* (Fox 2003), *MASS* (Ripley et al. 2013), *plyr* (Wickham 2011), *dplyr* (Wickham et al. 2019a), *devtools* (Wickham et al. 2019b), *ggbiplot* (Vu 2011), *psych* (Revelle 2018), *ggplot2* (Wickham & Chang 2008), and *outliers* (Komsta 2011).

## 3. RESULTS

### 3.1. Overview

Overall, the mud snail *Heleobia australis* showed a strong variation in snail density and a remarkable variation in several traits linked to contrasting biotic and abiotic pressures (environmental stress, predation, or parasite infection) occurring along the vertical zonation of the intertidal area. Tables 1 to 3 summarize descriptive statistics on density and shell traits measured for the first and fourth snail categories (cat. *i* and *iv*). In the appendix, we reported results of the snail density split by age classes (juvenile and adult; Table S1), principal components analyses (Text S3, Table S2, Fig. S3), descriptive statistics and the linear mixed models for the four snail categories on shell traits (Tables S3–S8) and for shell weight and body mass (Table S9).

**Table 1.**
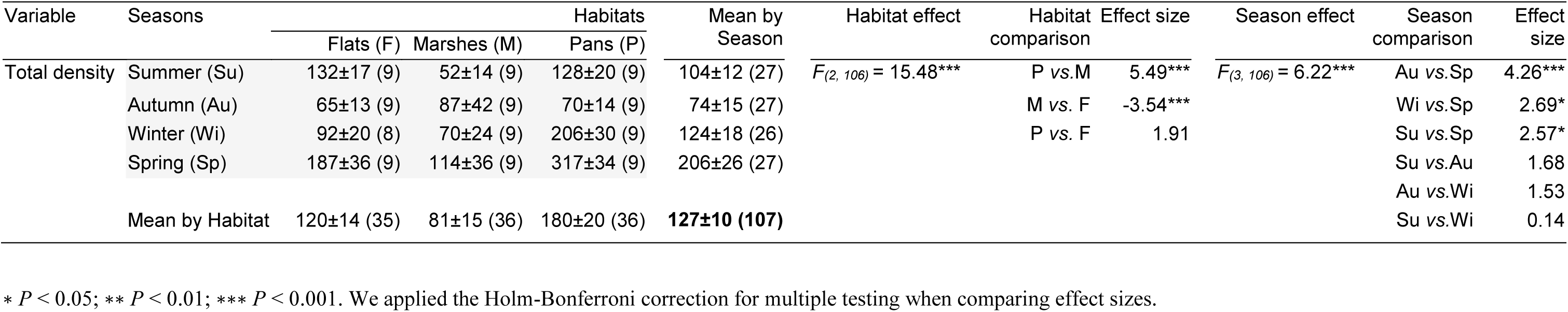
Summary of means (±SE) and statistical significance of the two-way ANOVA testing for habitat and seasonal effect on total snail density (ind./per sample; one sample being 78.5 cm^-2^) of the intertidal mud snail *Heleobia australis* from the Bahía Blanca estuary, Argentina. Number of samples are indicated between parentheses. Value in bold indicates overall mean. We reported raw estimates whereas the model fits for Ln-transformed density. See Methods section for details.

| Variable | Seasons | Habitats |  |  | Mean by Season | Habitat effect | Habitat comparison | Effect size | Season effect | Season comparison | Effect size |
| --- | --- | --- | --- | --- | --- | --- | --- | --- | --- | --- | --- |
|  |  | Flats (F) | Marshes (M) | Pans (P) |  |  |  |  |  |  |  |
| Total density | Summer (Su) | 132±17 (9) | 52±14 (9) | 128±20 (9) | 104±12 (27) | $F_{(2, 106)} = 15.48^{***}$ | P vs.M | 5.49<sup>***</sup> | $F_{(3, 106)} = 6.22^{***}$ | Au vs.Sp | 4.26<sup>***</sup> |
| | Autumn (Au) | 65±13 (9) | 87±42 (9) | 70±14 (9) | 74±15 (27) | | M vs. F | -3.54<sup>***</sup> | | Wi vs.Sp | 2.69* |
| | Winter (Wi) | 92±20 (8) | 70±24 (9) | 206±30 (9) | 124±18 (26) | | P vs. F | 1.91 | | Su vs.Sp | 2.57* |
| | Spring (Sp) | 187±36 (9) | 114±36 (9) | 317±34 (9) | 206±26 (27) | | | | | Su vs.Au | 1.68 |
| | Mean by Habitat | 120±14 (35) | 81±15 (36) | 180±20 (36) | **127±10 (107)** | | | | | Au vs.Wi | 1.53 |
|  |  |  |  |  |  |  |  |  |  | Su vs.Wi | 0.14 |

\* $P < 0.05$ ; \*\* $P < 0.01$ ; \*\*\* $P < 0.001$ . We applied the Holm-Bonferroni correction for multiple testing when comparing effect sizes.

**Table 2.**
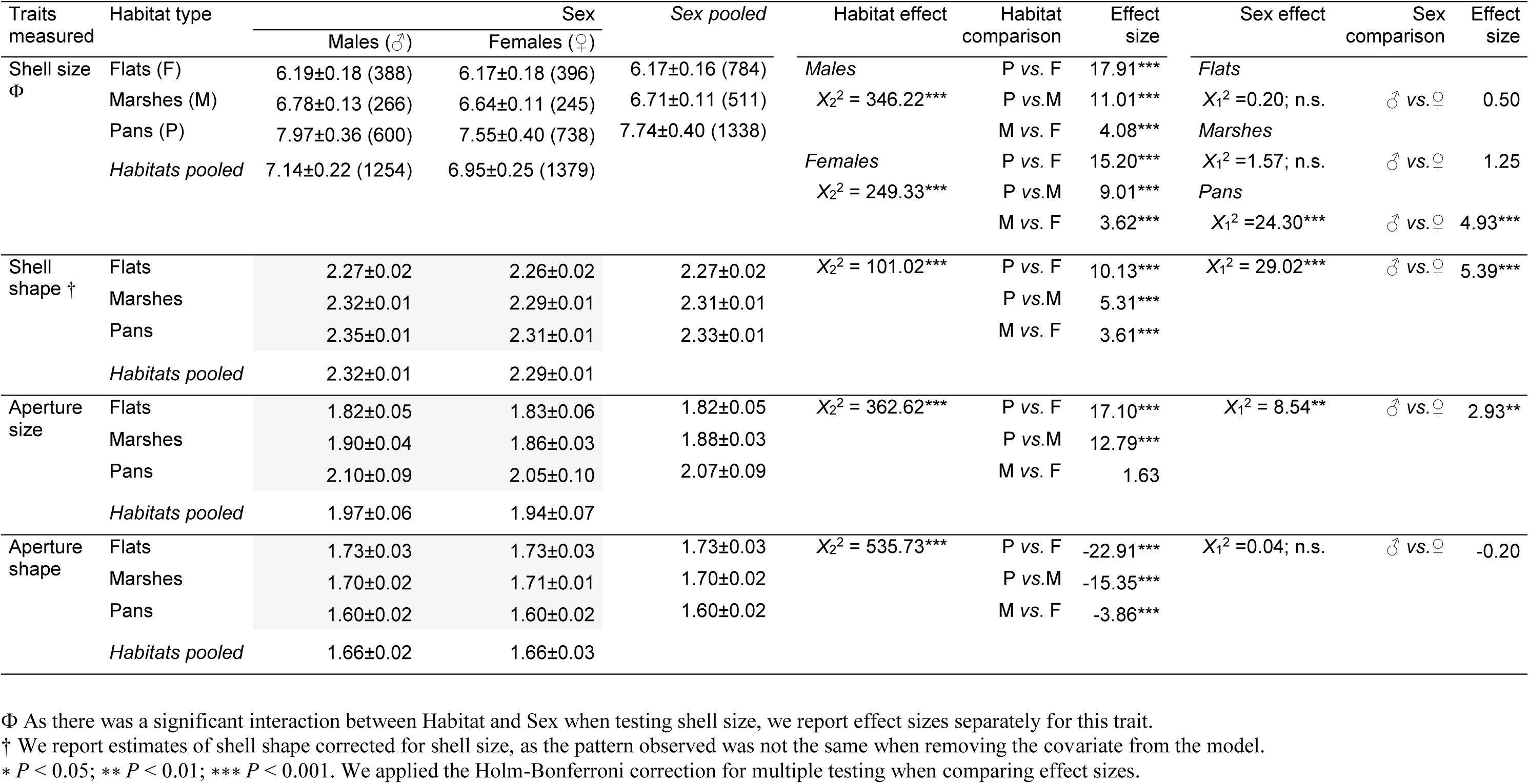
Uninfected adult snails. Summary of means (±SE) and statistical significance of habitat and sex effect on shell and aperture morphometrics of uninfected adult individuals of the mud snail *Heleobia australis* from the intertidal area of the Bahía Blanca estuary, Argentina. Variables are the same as indicated in table 2. Number of observations are indicated between parentheses, which are only shown for shell size but are the same for other traits measured.

**Table 3.** Infected vs. Uninfected adult snails. Summary of means (±SE) and statistical significance of status (infected of uninfected) and sex effect on shell and aperture morphometrics of the intertidal mud snail *Heleobia australis* from the Bahía Blanca estuary, Argentina. In these analyses we only considered individuals from pans, which allowed for preventing habitat effect on morphometry. Shell size estimated as the volume of a cone (mm^3^), shell shape as the length to width ratio (SL/SW), aperture size as the area of an ellipse (mm^2^), and aperture shape as the ratio between aperture length and width (AL/AW). Number of observations are indicated between parentheses, which are only shown for shell size but are the same for other traits measured.

| Traits measured | Habitat type | Sex |  | Sex pooled | Status effect | Status comparison | Effect size | Sex Effect | Sex comparison | Effect size |
| --- | --- | --- | --- | --- | --- | --- | --- | --- | --- | --- |
|  |  | Male (♂) | Female (♀) |  |  |  |  |  |  |  |
| Shell size | Uninfected (U) | 7.97±0.36 (600) | 7.55±0.40 (738) | 7.74±0.38 (1338) | $X_1^2 = 149.69^{***}$ | U vs.I | 12.42<sup>***</sup> | $X_1^2 = 31.68^{***}$ | ♂ vs.♀ | 5.65<sup>***</sup> |
|  | Infected (I) | 7.01±0.14 (432) | 6.76±0.20 (573) | 6.86±0.16 (1005) |  |  |  |  |  |  |
|  | Status pooled | 7.59±0.28 (1032) | 7.22±0.32 (1311) |  |  |  |  |  |  |  |
| Shell shape<br>Φ | Uninfected | 2.36±0.01 | 2.32±0.01 | 2.34±0.01 | Males | U vs.I | 3.43<sup>***</sup> | Uninfected | ♂ vs.♀ | 6.12<sup>***</sup> |
| | Infected | 2.40±0.01 | 2.39±0.02 | 2.40±0.01 | $X_1^2 = 11.82^{***}$ | | | $X_1^2 = 37.51^{***}$ | | |
|  | Status pooled | 2.38±0.01 | 2.35±0.01 |  |  |  |  |  |  |  |
|  |  |  |  |  | Females | U vs.I | 9.98<sup>***</sup> | Infected | ♂ vs.♀ | -0.30 |
| | | | | | $X_1^2 = 99.50^{***}$ | | | $X_1^2 = 0.09$; n.s. | | |
| Aperture size | Uninfected | 2.10±0.09 | 2.05±0.10 | 2.07±0.09 | $X_1^2 = 408.65^{***}$ | U vs.I | 21.11<sup>***</sup> | $X_1^2 = 14.95^{***}$ | ♂ vs.♀ | 3.88<sup>***</sup> |
|  | Infected | 1.80±0.04 | 1.76±0.06 | 1.77±0.05 |  |  |  |  |  |  |
|  | Status pooled | 1.98±0.07 | 1.92±0.08 |  |  |  |  |  |  |  |
| Aperture shape | Uninfected | 1.60±0.02 | 1.60±0.02 | 1.60±0.02 | $X_1^2 = 60.57^{***}$ | U vs.I | 7.83<sup>**</sup> | $X^2_1 = 1.3\text{e-}3$; n.s. | ♂ vs.♀ | 0.03 |
|  | Infected | 1.56±0.02 | 1.56±0.02 | 1.56±0.02 |  |  |  |  |  |  |
|  | Status pooled | 1.58±0.02 | 1.58±0.02 |  |  |  |  |  |  |  |

$\Phi$ As there was a significant interaction between Status and Sex when testing shell shape, we report effect sizes separately for this trait.
\* $P < 0.05$ ; \*\* $P < 0.01$ ; \*\*\* $P < 0.001$ . We applied the Holm-Bonferroni correction for multiple testing when comparing effect size

### 3.2. Strong habitat and seasonal effect on snail density

We found spatial differences in total density (including juveniles and adult snails; Table 1; Fig. 2). We observed no significant interaction between habitats and seasons (*F*_(6, 106)_ = 1.56, *P* = 0.167). Marshes showed the lowest density whereas flats and pans showed a marginal non-significant difference (Table 1). We found significant differences between seasons (Table 1). Density was higher in spring than in autumn, winter, and summer (Table 1). Snail density in autumn did not differ from that of summer or winter, nor did it differ significantly between summer and winter (Table 1), but it largely increased from autumn to spring (particularly in pans as compared to flats and marshes; Fig. 2). This increase across seasons was mainly driven by juvenile recruitment (Table S1; Fig. S2). By contrast, the effect of habitat conditions and seasons on adult density showed a significant interaction because density was higher in pans from autumn to winter whereas it decreased in marshes for the same period, and in flats it decreased from summer to autumn and then remained fairly constant (Table S1; Fig. S2).

**Figure 2.**
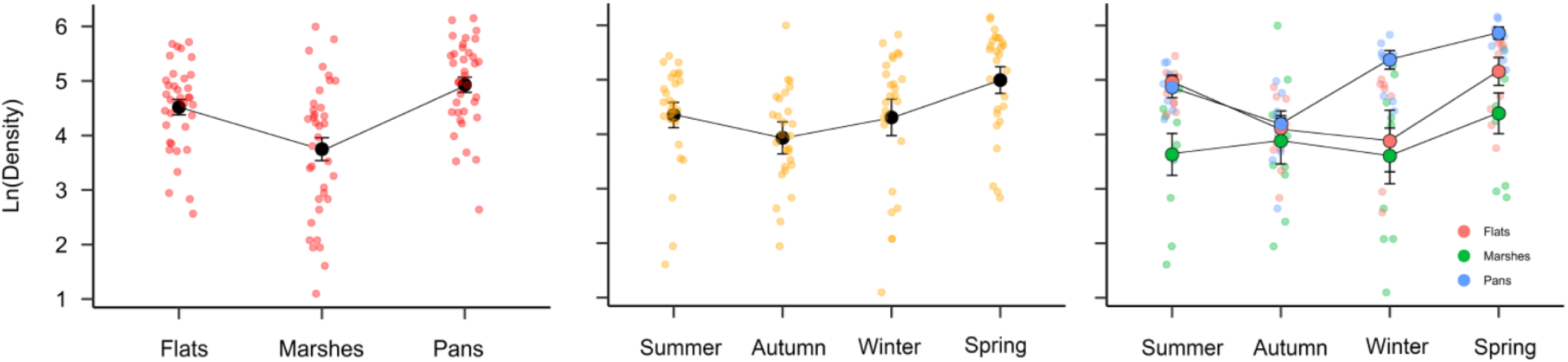
Variation in total density (juveniles and adults pooled) of the planktotrophic snail *Heleobia australis* across habitats and seasons in the intertidal area of the Bahía Blanca estuary, Argentina. Bars represent ± 1 SE.

**Figure 3.**
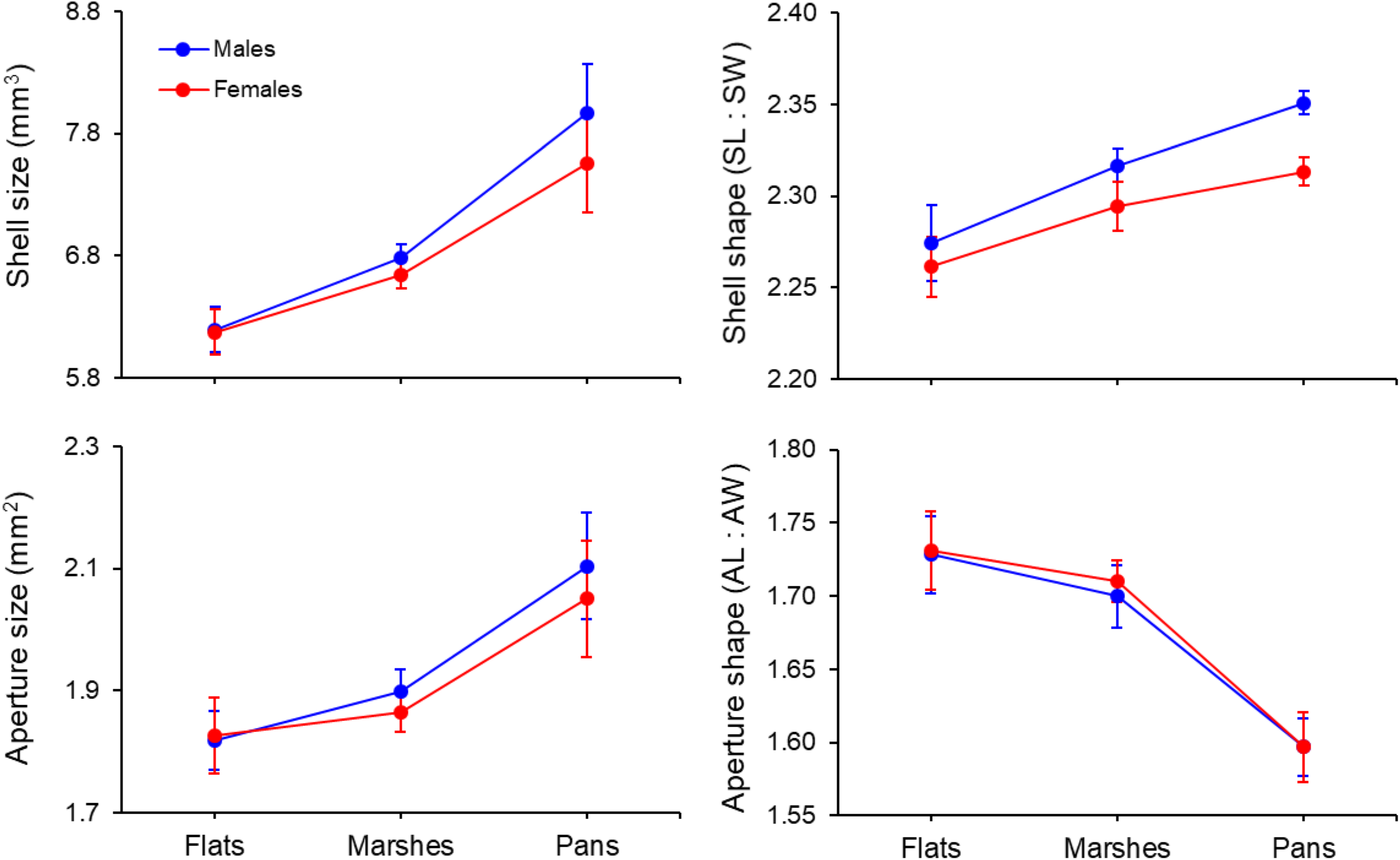
Phenotypic variation in shell and aperture morphometrics of uninfected adult snails *Heleobia australis* (cat. *i*) in the Bahía Blanca estuary, Argentina. Blue and red dots indicate mean values of each variable and sex in each habitat (flats, marshes, and pans). Mean values of shell shape were statistically corrected for shell size. Bars represent ± 1 SE.

### 3.3. Strong habitat effect on shell morphometrics

The morphometric responses to habitat conditions were similar in all three of the snail categories analyzed (cat. *i*–*iii*). For simplicity, we present results and figures only for uninfected adult snails (cat. *i*) in the main text and those for infected adults (cat. *ii*) and juvenile snails (cat. *iii*) are shown in the appendix (Tables S4-S7; Figs. S4-S5).

#### 3.3.1. Shell size

We found a significant interaction between fixed effects (habitat and sex; *X^2^* = 8.42, *P* = 0.015). This was mainly driven by shell size differences between sexes in pans, whereas such difference between males and females was not detected in snails from flats and marshes. In other words, males exhibited a larger shell size (mm^3^) compared to females in pans. This means that, in pans, the sex effect was much stronger in creating size differences between sexes compared to flats and marshes where the sex effect was weaker, which resulted in no difference in size between sexes. By contrast, the effect of habitat conditions was stronger in both sexes. Both male and female snails from flats and marshes showed a smaller shell size (a decrease of 12 and 22 % in shell size; respectively) relative to individuals from pans (Tables 2 and S3; Fig. 2).

#### 3.3.2. Shell shape

The interaction between fixed effects was significant (*X^2^_2_* = 8.20, *P* = 0.017), but when we statistically corrected for shell size, it became marginally non-significant (*X^2^_2_* = 5.69, *P* = 0.058). Snails from pans (sex pooled) exhibited the most elongated shells relative to individuals from marshes and flats. At the sex level (habitats pooled), males had a more elongated shell shape than females (Table 2 and S3; Fig. 2).

#### 3.3.3. Aperture size

We found a strong habitat and sex effect on aperture size; the interaction between these variables was not significant (*X^2^_2_* = 3.05, *P* = 0.217). Across habitats (sex pooled), snails from pans showed the largest aperture size (mm^2^) whereas in flats and marshes it was 9 to 12% smaller, respectively. Moreover, we observed a significant difference in aperture size between sexes (habitats pooled), males had bigger apertures than females and this difference was consistent across habitats (Table 2 and S3; Fig. 2).

#### 3.3.4. Aperture shape

We found no significant difference in aperture shape between sexes (*X^2^_1_* = 0.65, *P* = 0.724), and this pattern was similar even after correcting for shell size (*X^2^_2_* = 2.20, *P* = 0.333). However, individuals from pans (sex pooled) showed the most rounded shape relative to those from flats and marshes, which showed a more elongated aperture shape (Table 2 and S3; Fig. 2).

### 3.4. Pronounced trematode effect on shell morphometrics

Infection by the trematode *M*. *simillimus* strongly decreased mean values of most phenotypic traits of *H*. *australis* compared to uninfected snails from pans.

#### 3.4.1. Shell size

Shell size of infected snails (sex pooled) decreased 10% relative to uninfected ones, whereas females (infection status pooled) were 5% smaller than males (Table 3; Fig. 4). We found no significant interaction between the fixed effects (status and sex; *X^2^* = 0.70, *P* = 0.402).

**Figure 4.**
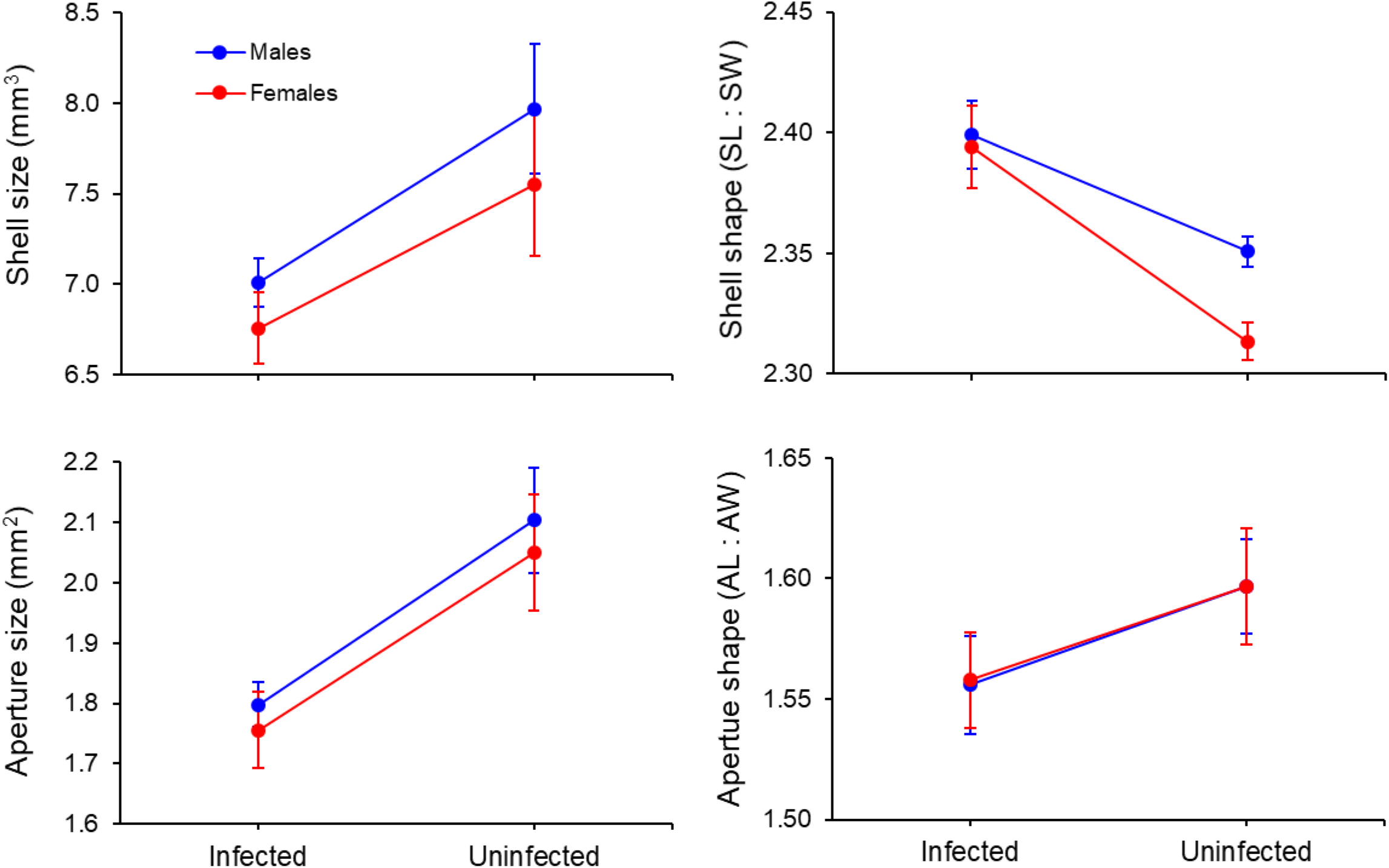
Phenotypic variation in shell and aperture morphometrics of infected and uninfected adult snails *Heleobia australis* (cat. *iv*) in response to a strong parasite pressure from a habitat with low environmental stress conditions but high parasite prevalence (pans) in the Bahía Blanca estuary, Argentina. Blue and red dots indicate mean values of each variable and sex (males and females, respectively). Mean values of aperture size were statistically corrected for shell size. Bars represent ± 1 SE.

#### 3.4.2. Shell shape

We found a significant interaction between status and sex (*X^2^* = 15.86, *P* <0.001). This was because, at the status level, uninfected males were more elongated (higher SL to SW ratio) than uninfected females, but infected males and females showed no difference in shell shape. We found the same pattern at the sex level; that is, infected male and female snails have a more elongated shell relative to uninfected male and female individuals (Table 3; Fig. 4).

#### 3.4.3. Aperture size

We observed a strong effect of infection status and sex in aperture size. Infected snails (sex pooled) and females (status pooled) had a smaller aperture than uninfected and male snails (15 and 3% respectively; Table 3; Fig. 4). We found no significant interaction between fixed effects (*X^2^_1_* = 2e-04, *P* = 0.990).

#### 3.4.4. Aperture shape

Uninfected snails exhibited a more elongate aperture shape (higher AL to AW ratio) than infected individuals whereas we found no sex effect on this shell trait (Table 3; Fig. 4). We found no significant interaction between fixed effects and no sex effect on aperture shape, even after correcting by shell size (*X^2^_1_* = 0.33, *P* = 0.564).

### 3.5. Thicker shells and lower body mass in habitats with high (a)biotic stress

We found that snails differed in shell weight (∼ thickness) across habitats (*X^2^_2_* = 48.93, *P* <0.001). Snails from pans showed the lightest/thinnest shells (11.53±0.23 mg) compared to snails from marshes (12.54±0.15 mg; *P*_Pans<Marshes_ <0.001) and flats (12.00±0.15 mg; *P*_Pans<Flats_ = 0.001). By contrast, individuals in marshes showed the heaviest/thickest shells (*P*_Marshes>Flats_ <0.001; Fig. 4). Body mass also differed across habitats (*X^2^_2_* = 19.67, *P* <0.001). Snails from pans exhibited the heaviest body mass (0.50±0.05 mg) relative to individuals from marshes (0.32±0.03 mg) and flats (0.33±0.04 mg) (*P*_Pans>Marshes_ <0.001; *P*_Pans>Flats_ <0.001), which showed no differences in body weight (*P*_Marshes<Flats_ = 0.390; Fig. 5). Further details on observations, corrected and uncorrected mean values, and statistics are indicated in the appendix (Text S4; Table S9; Fig. S6).

**Figure 5.**
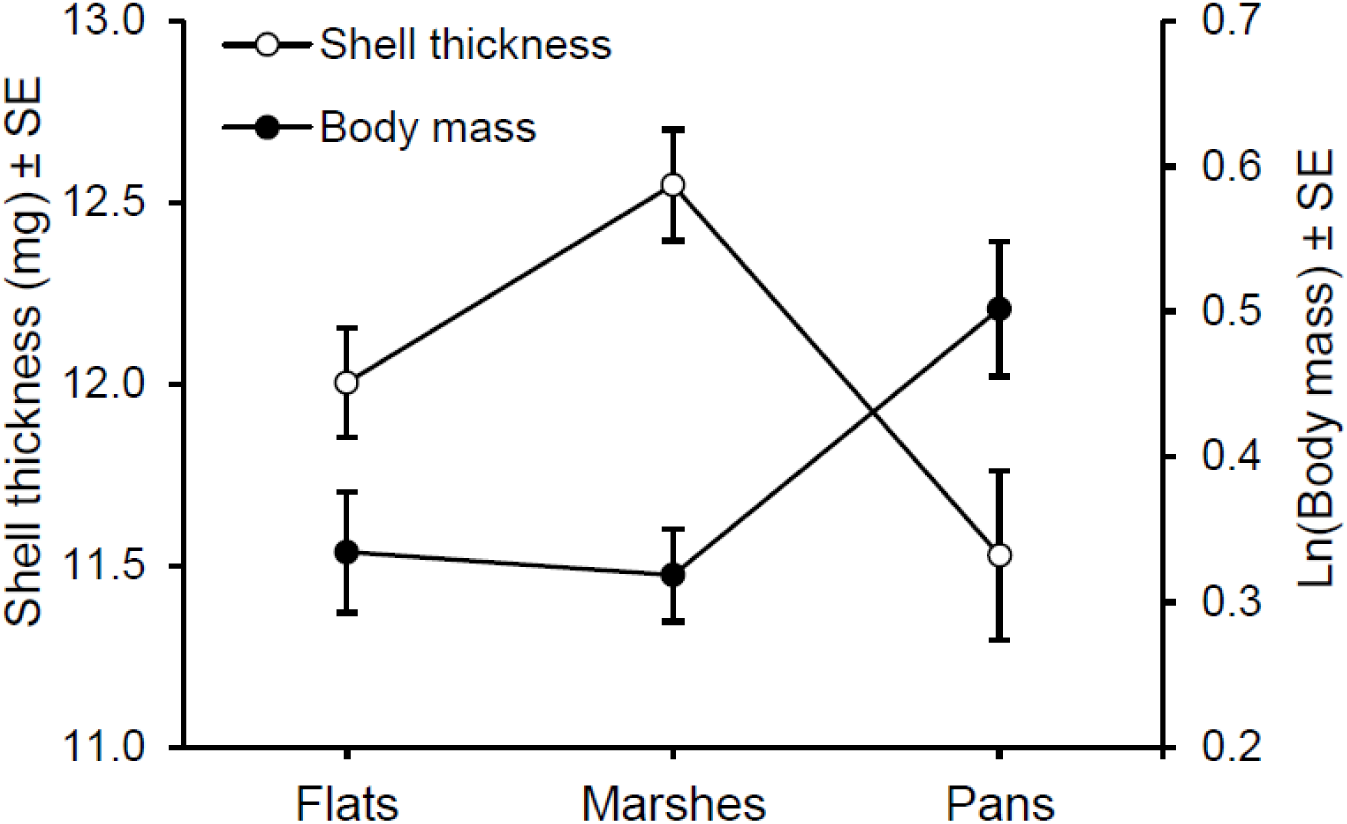
Shell weight (as proxy of thickness) and body mass variation of the intertidal mud snail *Heleobia australis* across habitats with different biotic and abiotic selective pressures in the Bahía Blanca estuary, Argentina. Bars represent ± 1 SE.

### 3.6. Variation in penis size

The penis shape confirmed that individuals from each habitat of the Bahía Blanca estuary belong to the same species (Fig. S8; Gaillard & Castellanos 1976). We found, however, a clear difference in penis size (*F*_(2, 39)_ = 8.42, *P* <0.001). Individuals from marshes showed the largest size relative to flats (*P*_flats<marshes_ <0.001) and pans (*P*_pans<marshes_ = 0.034); snails collected from these two habitats showed a marginal non-significant difference (*P*_flats<pans_ = 0.090; Fig. 6). We found no significant effect of shell size, meaning that differences in penis size were not due differences in individual’s shell size (*F*_(1, 39)_ = 0.27, *P*=0.608). Further, we found no differences in shell shape, even after controlling by shell size (*F*_(2, 43)_ = 0.851, *P* = 0.435).

**Figure 6.**
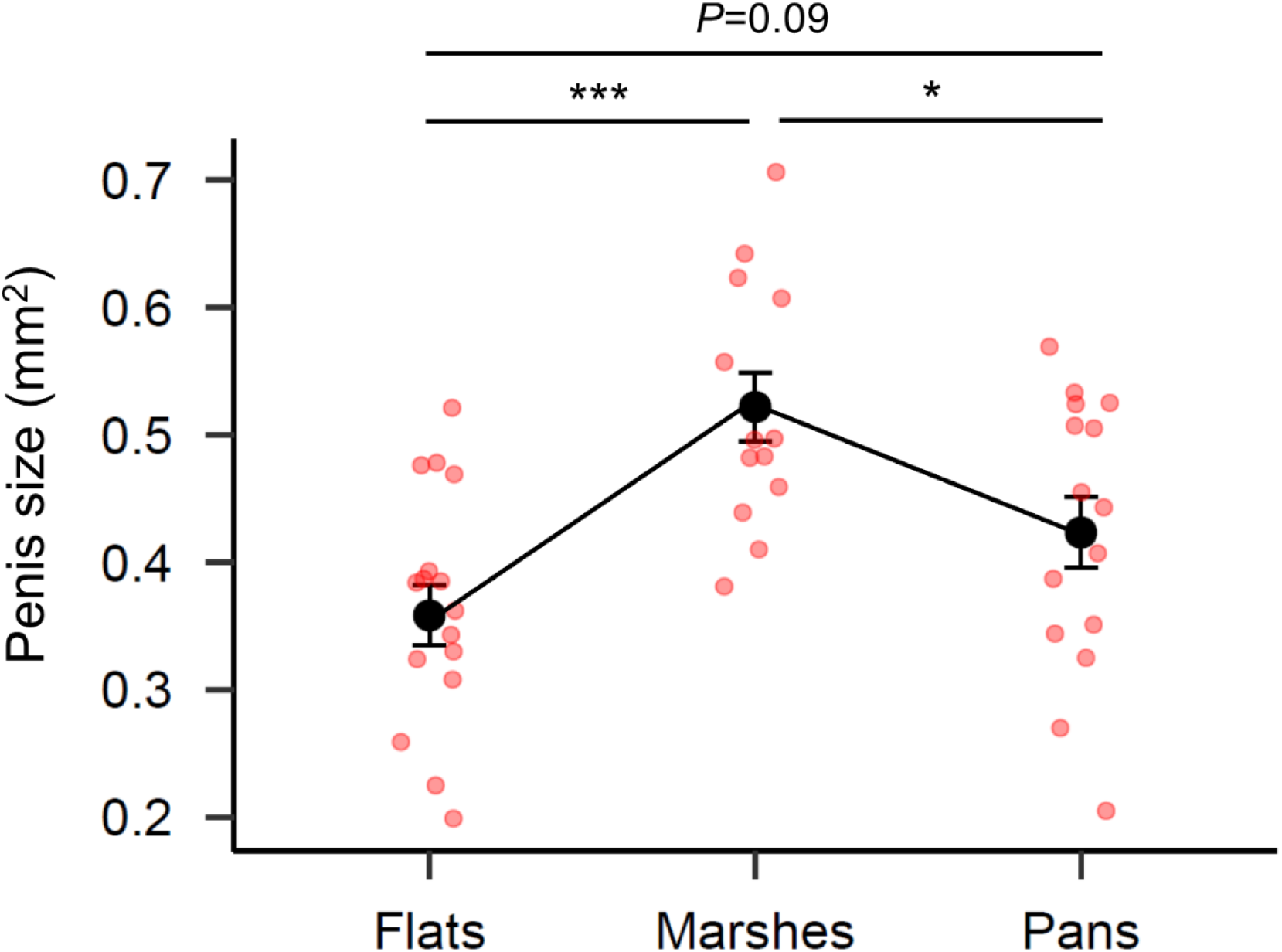
Differences in penis size (mm^2^) of uninfected adult snails *Heleobia australis* collected from three environmentally distinct habitats from the Bahía Blanca estuary, Argentina and kept in standard laboratory conditions for 21 months, which ensured that individuals analyzed were all adults. Bars represent ± 1 SE.

### 3.7. Molecular analysis of the cytochrome oxidase subunit 1 (COI) gene

The species-delimitation analysis implemented in ABGD found only one partition (prior maximal distance, *P* = 0.001), confirming that the individuals studied constitute a single *Heleobia* species. This finding is in agreement with the similar penis shape found among habitats. In fact, we did not observe any type of genetic structure among individuals. The gene tree and haplotype network showed that individuals did not gather in clusters depending on the habitat of origin (Figs. S8 and S9).

## 4. DISCUSSION

This study provides compelling evidence of ecological and phenotypic trait variation in response to contrasting biotic and abiotic conditions at a small spatial scale (summarized in Fig. 7). Our findings suggest that sympatric subpopulations of the intertidal snail *Heleobia australis* experienced a fine-grained environment. We hypothesize that the combined effect of phenotypic plasticity and local selection within each of the three habitats has favored the expression of potentially beneficial traits at a local scale.

**Figure 7.**
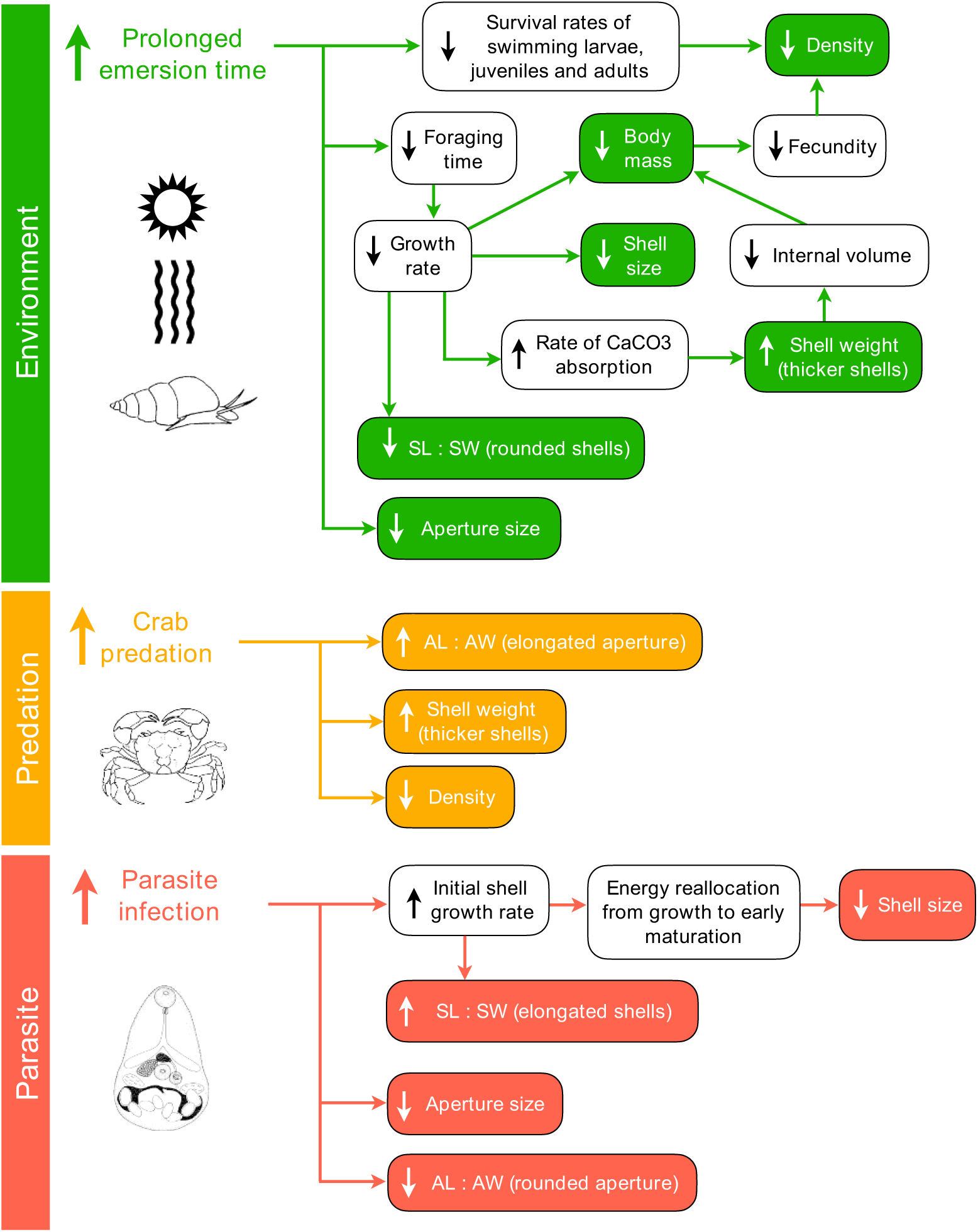
Summary of the responses observed in snail population density, shell and aperture morphometrics, and shell and body weight of the intertidal mud snail *Heleobia australis* from the Bahía Blanca estuary, Argentina. Such responses were induced by: (i) Environmental pressure, whereby snails experiencing prolonged emersion times in habitats that remain exposed to hot and dry conditions during low tide (marhses and flats) exhibited lower density and lower mean values of morphological traits relative to those from less environmental stress conditions (pans), which showed opposite phenotypic responses; (ii) predation pressure caused by the presence of the burrowing crab *Neohelice granulata* could be linked to the remarkable low density in marshes and could have favored the expression of shell defenses as shell thickening, increased shell globosity (lower SL to SW ratio), narrower apertures (higher AL to AW ratio); and (iii) strong parasite pressure caused by *Microphallus simillimus* where morphological traits of infected (I) snails shifted to lower mean values compared to uninfected (U) ones; note that phenotypic differences are only shown for pans, where parasite prevalence was the highest. Responses observed in this study are indicated by color-filled boxes, white-filled boxes indicate possible mechanisms or processes that might explain the observed responses. Green color refers to responses induced by environmental pressure, orange to phenotypic variation induced by crab predation, and pink by parasite pressure. Upper arrows (↑) indicate increase/higher/longer and down arrows (↓) indicate decrease/lower/shorter.

### 4.1. Reduced snail density under high environmental and predatory stress

Density of *Heleobia australis* was the lowest in marshes (vegetated area that drains at low tide) relative to flats (unvegetated area that drains at low tide) and pans (covered by water at low tide) whereas flats and pans showed no difference. However, when analyzed by age classes, adult density was higher in pans during winter but remarkably lower in marshes, even compared to flats (both habitats from the upper area). This is consistent with the idea that prolonged submersion times could increase the survival rate of snails during harsh climatic conditions (Fig. S2). As for juvenile snails, their density increased from autumn to winter in all the three habitats and kept increasing until spring, where it was 1.6 and 2.4 times higher in pans relative to flats and marshes, respectively (Table S1; Fig. S2). We found little support for the idea that intense environmental stress, combined with harsh climatic conditions, could strongly decrease their survival rate affecting their abundance, especially in the upper intertidal zone. Instead, these findings suggest that conditions at pans possibly favored juvenile recruitment, which could explain the differences observed in spring.

It is puzzling, however, that total density in marshes was lower relative to flats because *H. australis* is positively associated with marsh plants. Plants such as the smooth cordgrass *Spartina alterniflora*, which dominates the lower marsh, buffer physical stress factors (thermal and dehydration stress) relative to uncovered areas, promoting snail aggregation <u>(Canepuccia et al. 2007</u> and references therein). One possible explanation for the low density in marshes, relative to flats, is a possible negative interaction between *H*. *australis* and the grapsid crab *Neohelice granulata*, whose aggregation is also facilitated by marsh plants (Spivak et al. 1994, Alvarez et al. 2013, Angeletti et al. 2014, Angeletti & Cervellini 2015). The crab attains density peaks in late spring and early summer but is absent throughout the year in unvegetated areas (flats or pans; (Angeletti & Cervellini 2015). It is therefore likely that the low density of *H*. *australis* in vegetated crab-rich habitats resulted from an increased bioturbation activity caused by *N. granulata*, particularly during spring time (Alvarez et al. 2013). Another mutually non-exclusive hypothesis is that this difference between marsh and flat density could result from a higher predation pressure caused by this crab (D’Incao et al. 1990, Barutot et al. 2011), in which case it might have induced predatory shell defenses such as shell thickening or narrower apertures (Boulding & Hay 1993, Johannesson & Johannesson 1996, Rolán-Alvarez 2007, review by Bourdeau et al. 2015). Snails from marshes exhibited a strong variation in shell characters that supports this hypothesis, which we discuss later in this section.

### 4.2. Snails exhibited larger, narrower, and thinner shells and heavier body mass in habitats with low stress conditions

Snails subjected to different levels of stress showed a strong phenotypic variation. In all categories analyzed (cat. *i*–*iii*), individuals showed a clear shift in their phenotype, likely in response to different environmental pressures (Fig. 7). Overall, the distribution of trait values tended to represent two distinct groups, one for flats and marshes and another for pans. Such responses seem to be primarily linked to the level of exposure to physical stress (temperature and salinity fluctuation and desiccation) of individuals during low tide. Prolonged submersion times enhance foraging times and the absorption of calcium carbonate from water increases shell growth rate, which in turn gives rise to larger but more elongated shells (review by Chapman 1995). As there is a maximal rate at which calcium carbonate can be absorbed from the water, rapidly growing individuals produce thinner shells for the same amount of body weight than slower growing individuals. This results in a larger internal volume that accommodates a higher body mass and is related to higher growth rate favored by longer foraging times (Palmer 1981, Kemp & Bertness 1984, Chapman 1995, Trussell 2000a). Accordingly, snails that remain covered by water during low tide (pans) showed larger, narrower, and thinner shells and heavier body mass relative to individuals from habitats with high environmental stress conditions (flats and marshes). As higher body weight is directly linked to higher fecundity (i.e. egg production; Fredensborg et al. 2006), it is therefore likely that the heaviest body weight of individuals from pans was related to a higher recruitment, which would explain the increased juvenile abundance compared to flats and marshes (Fig. S2).

Juvenile and adult snails from flats and marshes showed a smaller aperture size (smaller aperture surface area) with respect to individuals from pans. It could be that higher environmental stress conditions favored the expression of a smaller aperture size as a potentially beneficial trait, thus increasing resistance to high desiccation in these habitats that drain at low tide. This is consistent with other studies showing that the size of the shell aperture in gastropods is smaller in response to hot and dry conditions (Machin 1967, Vermeij 1973, Chapman 1995, Melatunan et al. 2013, Schweizer et al. 2019). Moreover, individuals from flats and marshes showed thicker shells and lower body mass. Shell thickening could be due to a higher deposition of calcium carbonate at a slow grow rate (likely in response to unfavorable conditions in more exposed habitats from the upper intertidal) that traded off against investment in body mass (Palmer 1981).

### 4.3. Crab predation pressure would favor the expression of shell defenses in *Heleobia australis*

Snails from marshes showed a more rounded and thicker shell, reduced body mass, and a narrow aperture shape for three snail categories analyzed (cat. *i*–*iii*; Fig. 7). Such characteristics are consistent with shell defenses probably induced by the high presence of the predatory crab *Neohelice granulata* in that particular habitat (e.g. Appleton & Palmer 1988, Palmer 1990, Bourdeau et al. 2015). Considering that crushing predators exert a strong selective pressure driving the evolution of behavioral, chemical, and morphological defense traits (Appleton & Palmer 1988, Palmer 1990, Trussell 2000b, Johannesson 2003, Bourdeau et al. 2015), our results, combined with the pattern of remarkably low density, support the hypothesis that predation pressure is stronger in marshes than in flats and pans. Yet, specific information on whether crabs actively predate on *H. australis* is limited or inexistent. Earlier studies reported that *N. granulata* consumes mollusks but in a low frequency (D’Incao et al. 1990, Barutot et al. 2011). These studies, however, have used visual examination of gut/stomach contents. This procedure may have under-estimated true predation levels as crabs only eat the soft tissue after crushing/peeling the shell, whereas the remains found in guts or stomach become unidentifiable due to maceration and digestion. Thus, further studies using molecular-based tools are needed for detecting the presence of *Heleobia* tissue in crab stomachs as an alternative or complementary approach to visual identification (e.g. Albaina et al. 2010, Collier et al. 2014). This would establish the trophic link between these two species that is highly likely to exist. Evidence from this study clearly reveals that *H*. *australis* phenotypically responded to the presence of predatory crabs, even early in life as juveniles from marshes also showed a narrow aperture shape and a more rounded shell. Our results therefore warrant further investigation to understand the adaptive value of the plastic shell responses of gastropods induced by predators.

### 4.4. Strong parasite effect on shell characters: a trade-off between growth and early reproduction

Infected and uninfected snails from pans showed a clear phenotypic differentiation in shell traits. Trematode infection inevitably leads to snail host castration and drastically reduces host fitness, which, in evolutionary terms, is equivalent to death of the host (Lafferty 1993, Fredensborg et al. 2006). In this sense, being castrated by a parasite is similar to being eaten by a predator (Kuris 1974). This means that parasitic castrators can exert a strong selective force that could favor the expression of behavioral, physiological, and morphological adaptations to minimize the negative impact of parasitism on host fitness (Lafferty 1993). Early maturation, which results from fast growing individuals, is an effective strategy that increases current reproductive effort over future reproduction (Cole 1954, Lewontin 1965, Roff 1992), which equals to 0 in castrated hosts. In *Heleobia*, castration caused by *Microphallus* can have a profound impact on snail fitness as this parasite shows an extensive host exploitation occupying the entire gonad and most of digestive gland (see Fig. S2 in Alda et al. 2019).

Our results showed that infected snails had a more elongated shell shape but a smaller size compared to uninfected snails (Fig. 7). Although infected snails were smaller, they exhibited the same number of whorls (six) as uninfected snails, which indicates that individuals analyzed were adults (Gaillard & Castellanos 1976). It could therefore be that the elongated shape resulted from an initial higher growth rate (review by Chapman, 1995) likely followed by an energy reallocation from growth to early reproduction (Lafferty 1993, Agnew et al. 1999), which could explain the smaller shell size compared to uninfected snails (Alda et al. 2010, 2019, this study). Infected snails also showed a smaller aperture size but a more rounded shape relative to uninfected individuals (Fig. 7). This change in aperture size and shape could be a side effect of selection for high rate of shell growth (Boulding & Hay 1993) induced by parasites. Together, these findings support the idea that infection by trematodes exerted a strong selective pressure on the snail host, causing important shifts in the expression of shell traits towards lower mean values, which created a pronounced phenotypic difference between infected and uninfected snails (Poulin & Thomas 1999).

### 4.5. Difference in penis size in a single *Heleobia* species

Interestingly, individuals from each habitat showed a clear variation in penis size in spite of their shared rearing in common garden laboratory conditions (no habitat effect). After dissection, we observed that none were infected by parasites (no parasite effect). Field-collected juveniles might retain the influence of environmental conditions experienced prior to collection (Sanford & Kelly 2011). Thus, one possible explanation is that this variation in penis size resulted from different growth rates as well as the differences in shell shape observed for wild snails. We found, however, no differences in shell shape among male individuals reared in laboratory conditions, even after controlling by shell size. It is noteworthy that a strong difference in penis size emerged despite the relatively small number of male individuals analyzed (n = 15 per habitat). One interpretation of this phenotypic divergence is that it was not related to dissimilar rates of growth but to genetic differences, suggesting that this is likely a non-plastic trait. Despite the difference in size, its shape indicates that individuals from each subpopulation belong to the same species, more specifically to *Heleobia australis australis* (see Gaillard & Castellanos, 1976 for further detail on taxonomy). Accordingly, the COI gene confirmed that specimens considered in this study constitute a single species.

Future experimental (laboratory breeding) and non-experimental approaches (multilocus or genomic analyses) are needed to exclude other processes that may have contributed to this difference in penis size. These approaches will help us to understand whether this genital divergence could be an indicative of pre-zygotic reproductive isolation among subpopulations of *Heleobia australis* (Kameda et al. 2009, Hollander et al. 2013). If this is true, it could be that individuals studied herein would be part of a species complex that have recently diverged, which would be in line with the lack of clusters when analyzing the conserved COI gene (Kemppainen et al. 2009, Janzen et al. 2017, Matos-Maraví et al. 2019).

### 4.6. Does the small-scale phenotypic differentiation in *Heleobia australis* reflect adaptive plasticity responses to local environmental pressures?

Theory predicts that adaptive phenotypic plasticity is more common than genetic adaptation in wide-dispersing species relative to species with restricted dispersal (Mather 1955, Levins 1968, Via & Lande 1985, Scheiner 1998, De Jong 1999, 2005). That is because gene flow homogenizes genotype frequencies across space, countering the effect of natural selection (review by Lenormand 2002). Our results suggest that sympatric (or micro-parapatric; Butlin et al. 2008) subpopulations of *Heleobia australis* experienced a fine-grained environment where conditions imposed by contrasting biotic and abiotic stressors across the vertical gradient favored the expression of beneficial traits. This study was not designed to test for an adaptive basis to phenotypic trait variation, but provides a reasonable ground for advocating that the shifts in the mean trait values could be adaptations in response to the prevailing selection pressure at each habitat. If this is true, adaptive plasticity might have placed populations close enough to the new favored phenotypic optimum for directional selection to act, thus foreshadowing adaptive evolutionary responses over longer periods of time (Ghalambor et al. 2007).

The question that inevitably arises is whether there is scope for local adaptation in this intertidal mud snail. An increasing number of experimental studies indicates that local genetic adaptation may be more common than predicted by theory in marine species with planktonic dispersal, especially in systems characterized by strong selection over relatively fine spatial scale (Hays 2007, Marshall et al. 2010, Sanford & Kelly 2011). One possible explanation for the evolution of locally adapted phenotypes in species with planktonic dispersal is linked to pre- and post-colonization barriers to connectivity (e.g. mortality due to predation, starvation, or environmental stress) that increase localized recruitment (i.e. phenotype-environment mismatch; Levin et al. 2006, Marshall et al. 2010). That is, local adaptation may be ensured by genetic polymorphism because immigration is rare and the intrapopulation gene pool is highly reliant upon local recruitment, where individuals are exposed to local selection pressures. Directional selection of adaptive traits may induce rapid morphological and ecological changes favoring genotypes exhibiting the most extreme phenotypes. This selection process would then alter the reaction norm allowing the population to reach the fitness optimum, whereby the induced phenotype becomes later genetically ‘assimilated’ (reviewed in Waddington 1953, West-Eberhard 1989, Pigliucci et al. 2006, but see West-Eberhard 2003). Only a few studies reported local genetic differentiation in wide-dispersing marine snails, contributing to phenotypic differentiation as a result of local disruption to dispersal of planktotrophic larvae, post-veligers, and juvenile stages (e.g. Struhsaker 1968, Parsons 1997). In this context, it is therefore plausible that the planktotrophic snail *H*. *australis* benefits from the combined effect of plasticity and directional selection to evolve locally adapted phenotypes to contrasting habitat conditions across the intertidal gradient.

## 5. CONCLUSION

Overall, our results showed that the same phenotype was unlikely to perform best regardless of the stress encountered. Contrasting environmental stressors across the vertical gradient induced important shifts in the expression of phenotypic traits, thus creating pronounced small-scale phenotypic differences. These findings support the perspective that phenotypic plasticity favored the expression of beneficial traits thereby increasing the likelihood of persistence of subpopulations facing local environmentally stressful conditions. The next step is therefore to perform more powerful experimental approaches (including common garden designs and reciprocal transplants) and molecular analyses to unveil the adaptive value of the small-scale phenotypic plasticity in this marine snail. Future studies are clearly needed to understand the spatial scale at which adaptive differentiation can occur and the relative contributions of phenotypic plasticity versus fixed genetic variation in key traits of wide-dispersing marine organisms. Such information is critical for improving the prediction of the biological impacts of climate change and for effective conservation management approaches of marine system populations.

## ACKNOWLEDGEMENTS

We would like to express our gratitude to Jelena Pantel, Kevin Lafferty, Patrice David, Antonio A. Vazquez Perera, Philippe Jarne, and Romain Villoutreix for their constructive suggestions on earlier versions of the manuscript. We also thank Leandro A. Hünicken and Micaela Folino for field and laboratory assistance and to Néstor J. Cazzaniga for providing insightful comments on the systematics of *Heleobia* group. This manuscript was improved by comments from Dr. Sokolova and three anonymous reviewers. This work was partially funded by a research grant from the Universidad Nacional del Sur (UNS 24/B153 and UNS 24/B199). NB was partially supported by the “Programa de Financiamiento Parcial de Estadías en el Exterior para Investigadores Asistentes”, National Scientific and Technical Research Council CONICET (Res. N° 558 1236/08; 4118/16)”.

## AUTHORS’ CONTRIBUTIONS

NB and PA conceived the ideas and designed methodology; NB, JPP, and PA collected the data; NB analyzed the data and led the writing of the manuscript. All authors contributed critically to the drafts and gave final approval for publication.

