## Appendix for "Environmental stressors induced strong small-scale phenotypic differentiation in a wide-dispersing marine snail"

### Table of Contents

***Supplementary Methods and Results***
*Text S1.* Size frequency distribution and cohort identification 2
*Text S2.* Principal components 2
*Text S3.* Strong habitat effect on shell morphometric 2
*Text S4.* COI analysis 3
References 4

***Supplementary Tables***
*Table S1.* Results of two-way ANOVAs for adult and juveniles snail density 6
*Table S2.* Results of PCAs 7
*Table S3.* Results of linear models for uninfected adults 8
*Table S4.* Descriptive statistics for infected adults 9
*Table S5.* Results of linear models for infected adults 10
*Table S6.* Descriptive statistics for uninfected juveniles 11
*Table S7.* Results of linear models for uninfected juveniles 12
*Table S8.* Results of linear models for infected and uninfected adults 13
*Table S9.* Results of linear models for shell and body weight 14

***Supplementary Figures***
*Fig. S1.* Length-frequency distributions and polymodal decomposition 15
*Fig. S2.* Spatial and seasonal variation in snail density 16
*Fig. S3.* Principal components analyses of shell morphometric 17
*Fig. S4.* Shell polymorphism of infected adult snails 18
*Fig. S5.* Shell polymorphism of juvenile snails 19
*Fig. S6.* Shell weight and body mass across habitat conditions 20
*Fig. S7.* Penis morphology 21
*Fig. S8.* Gene tree 22
*Fig. S9.* Haplotype network 23

### Supplementary Methods and Results

**Text S1. Size frequency distribution and cohort identification**
The length-class interval was calculated using the Sturges's method (Bonel, Solari, & Lorda, 2013). We then applied Bhattacharya's method available in FISAT II software (Version 1.2.0, FAO-ICLARM Fish Assessment Tools; Gayanilo, Sparre, & Pauly, 2002). To confirm each modal progression, we used the NORMSEP method also available in the FISAT II software (Pauly & Caddy, 1985). Those individuals outside the lower 95% confidence limit of the adult cohort were considered juveniles (Fig. S1).

**Text S2. Main components of the morphological shell variation within each snail group**
For each of the snail categories (cat. *i*–*iv*), principal component analyses decomposed the shell morphometric measures (shell and apertures size and shape) into four principal components of which the first two explained, overall, more than 73% of the variance in the original data (Table S2; Fig. S3). The component loadings showed that the shell and aperture size are highly correlated one another and with PC1, whereas PC2 highly contrast shell shape with aperture shape. In other words, the first component was strongly correlated with size indices, and the second component with shape (further details in Table S2). The distribution of individual data points from flats and marshes highly overlapped, and this was consistent in all the three snail categories (cat. *i*–*iii*). This constituted one group that clearly differed from the other group of data points that was composed by individual observations from pans (Fig. S3). Likewise, the scatter of the data showed two groups when decomposing the variation of morphometric measures between infected and uninfected adult snails (cat. *iv*).

**Text S3. Strong habitat effect on shell morphometric**
Infected adult snails
*Shell size*. Snails (sex pooled) from more stressed habitats showed a 4 to 10% decrease in shell size relative to snails from pans; there was no difference in size between snails from flats and marshes. Although the trend was similar to that of uninfected snails, the habitat effect in this group was less strong (Table S5-S6; Fig. S4). We also observed that males were bigger than females irrespective of their habitat, as indicated by the non-significant interaction between habitat and sex ($X^2_2$ = 0.43, $P$ = 0.807). *Shell shape*. Infected snails from pans showed the most elongated shell shape relative to those from flats and marshes, but unlike uninfected snails, we found no differences in shell shape between these two habitats nor between sexes, even though we corrected for shell size. We thus reported uncorrected estimates (Table S5-S6; Fig. S4). In addition, we found no significant interaction between fixed effects ($X^2_2$ = 2.66, $P$ = 0.265). *Aperture size*. This shell trait followed a similar trend as shell size. That is, snails from flats and marshes, irrespective of their sex, showed a 5% decrease in aperture size with respect to snails from pans. Males had a larger aperture than females and this pattern was observed across all the three types of habitats (Table S5-S6; Fig. S4), which is reflected by the non-significant interaction between categorical variables ($X^2_2$ = 1.22, $P$ = 0.542). *Aperture shape*. Differences across habitats mirrored the pattern observed for uninfected snails, meaning that individuals from marshes showed the most elongated aperture shape with respect to snails from flats and pans, which showed no differences (Table S5-S6; Fig. S4). We found no significant sex effect ($X^2_1$ = 5e-04, $P$ = 0.982) and no significant interaction ($X^2_2$ = 0.92, $P$ = 0.631). As we obtained a similar result after correcting for shell size, we therefore reported uncorrected estimates.

Juvenile snails

*Shell size*. We found a significant interaction between habitat and sex effects ($X^2_2$ = 8.22, $P$ = 0.016). This was because shell size in males was 46, 43, and 17% larger than females in flats, marshes, and pans; respectively. These figures show that sex effect was stronger in habitats with more stressful conditions than in those with low stressful conditions, which is similar to what we observed for uninfected adults (Table S7-S8; Fig. S5). *Shell shape*. There was no interaction between habitat and sex effects ($X^2_2$ = 1.48, $P$ = 0.477) even after correcting for shell size ($X^2_2$ = 0.01, $P$ = 0.994). Juvenile snails from pans (sex pooled) showed a more elongated shell shape than those from flats and marshes, which exhibited a more rounded shape. Males had a more elongated shape than females and it was consistent across habitats, and the same pattern was observed after correcting for shell size, so we reported uncorrected estimates (Table S7-S8, Fig. S5). *Aperture size*. This trait followed the same trend as shell size. The interaction term was significant ($X^2_2$ = 9.93, $P$ = 0.007) because the effect size between males and females from pans was smaller compared to that of flats and marshes (Table S7-S8; Fig. S5). On the one hand, juvenile males from habitats with high stress conditions showed, on average, a 7-8% decrease in aperture size relative to the males from low stress conditions. The habitat effect in females was much stronger representing a decrease of 22-23% in aperture size when comparing habitats with low vs. high stress conditions. We found no differences in aperture size neither for males nor females from flats and marshes. On the other hand, sex effect was stronger in more stressful habitat conditions where females showed a 31% decrease of aperture size with respect to males (both in flats and marshes), whereas in pans females had an aperture size 17% smaller than males (Table S7-S8; Fig. S5). *Aperture shape*. Habitat and sex effects showed no significant interaction ($X^2_2$ = 1.89, $P$ = 0.389). We obtained a similar result when we added shell size as covariate ($X^2_2$ = 1.70, $P$ = 0.426), but sex effect became non-significant after correction. So, we reported corrected estimates (Table S7-S8, S5). We found, however, a strong habitat effect that mirrored patterns observed for adults (both infected and uninfected), meaning that juveniles from flats and marshes showed a more elongated aperture shape whereas snails from pans exhibited a rounded shape (Table S7-S8; Fig. S5).

**Text S4. COI analysis**

To verify whether the individuals living in different habitats belong to the same species or to a species complex, we conducted molecular analyses by amplifying the cytochrome oxidase subunit 1 (COI) gene in 10 individuals sampled from each of the three sites. We used the primers LCOI490 (forward) 5’ GGTCAACAAATCATAAAGATATTGG 3’ and HCO2198 (reverse) 5’ TAAACTTCAGGGTGACCAAAAAATCA 3’ to amplify COI (Folmer et al. 1994). PCR amplification was performed in a total volume of 25 µl containing 12.5 µl of Taq PCR Master Mix (Qiagen), 2.5 µl of each primer (10 mM) and 2 µl of DNA in an Eppendorf Thermal Cycler with an initial denaturation step at 95 °C for 15 minutes; followed by 35 cycles of denaturation at 95 °C for 30 seconds, annealing at 50 °C for one minute, extension at 72 °C for one minute; and a final elongation at 60 °C for 30 minutes. The presence and size of amplification products were electrophoretically confirmed in 1% agarose gels stained with EZ-Vision. DNA sequencing was performed by Eurofins Genomics (Ebersberg, Germany) using PCR-amplified products as templates. All sequences were uploaded to GenBank (MT294016, MT295108-MT295136).

We searched the literature and GenBank for sequence data attributable to *Heleobia australis*. We found only one sequence which was included in our dataset: JQ972708 from Mar Chiquita coastal lagoon (Argentina)—some 600-km northwest from the Bahía Blanca estuary. Alignment was performed using MAFFT (Katoh & Standley, 2013). Ambiguously aligned sites were excluded using GBLOCKS with default settings for a less stringent selection (Castresana, 2000). The number of positions in the final sequences was 593 (87% of the original 677 positions).

We used Bayesian inference in Beast2 (Bouckaert et al., 2014). The best-fitting model of sequence evolution was selected using bModelTest (Bouckaert et al., 2014). The best model was HKY+G+I. The analyses were run using four gamma categories and a proportion of 0.5 invariant sites. We partitioned codon positions, unlinked substitution models and clocks and linked trees. A strict-clock model was chosen. We used a birth-death model as prior with lognormal birth and death rates. All the MCMC were run for 100,000,000 generations storing every 10,000 generations. The MCMC output was visualized using Tracer (Rambaut, Drummond, Xie, Baele, & Suchard, 2018) and tree samples summarized by TreeAnnotator (utility program distributed with the Beast package) using a 10% burn-in. The species tree was visualized and edited in FigTree and GIMP (https://www.gimp.org). We also built a TCS haplotype network using popART (Leigh & Bryant, 2015) and compared it with the gene tree.

We applied a species-delimitation method: Automatic Barcode Gap Detection (ABGD; (Puillandre, Lambert, Brouillet, & Achaz, 2012) using the default settings (https://bioinfo.mnhn.fr/abi/public/abgd/). ABGD does not rely on tree shapes but on divergence. The method detects barcode gaps, which can be observed whenever the divergence among organisms belonging to the same species is smaller than divergence among organisms from different species (Puillandre et al., 2012).

### SUPPLEMENTARY TABLES

**Table S1.** Summary of means (±SE) and statistical significance of the two-way ANOVAs testing for habitat and seasonal effect on adult and juvenile snail density (ind./per sample; one sample being 78.5 $cm^{-2}$) of the intertidal mud snail *Heleobia australis* from the Bahía Blanca estuary, Argentina. Number of samples are indicated between parentheses. We reported raw estimates whereas the models fits for transformed variables. See Methods section for details. We found a significant interaction between habitat and season effects when analyzing density of adult snails ($F_{(6, 106)}$ = 2.43, $P$=0.031).

| Variables | Seasons | Habitats | | | *Seasons* | Habitat effect | Habitat | Effect | Season effect | Season | Effect |
|---|---|---|---|---|---|---|---|---|---|---|---|
| | | Flats (F) | Marshes (M) | Pans (P) | *pooled* | | comparison | size | | comparison | size |
| Adults Φ | Summer (Su) | 100±11 (9) | 43±12 (9) | 116±17 (9) | 72±12(27) | *Summer* | | | *Flats* | | |
| | Autumn (Au) | 44±8 (9) | 62±28 (9) | 63±13 (9) | 57±10 (27) | $F_{(2, 27)}$ = 8.97** | P *vs.* F | 0.09 | $F_{(3, 35)}$ = 4.05* | Wi *vs.* Sp | -3e04 |
| | Winter (Wi) | 59±12 (8) | 27±8 (9) | 177±25 (9) | 89±16 (26) | | M *vs* . F | -1.18** | | Au *vs.* Wi | -0.19 |
| | Spring (Sp) | 53±9 (9) | 25±8 (9) | 104±13 (9) | 61±9 (27) | | P *vs.* M | 1.27** | | Au *vs.* Sp | -0.19 |
| | | | | | | *Autumn* | | | | Su *vs.* Wi | 0.70 |
| | *Habitats pooled* | 73±11 (35) | 72±11 (36) | 76±10 (36) | | $F_{(2, 27)}$ = 0.62; n.s. | — | — | | Su *vs.* Sp | 0.70 |
| | | | | | | *Winter* | | | | Su *vs.* Au | 0.88* |
| | | | | | | $F_{(2, 27)}$ = 12.15*** | M *vs* . F | -1.19 | *Marshes* | | |
| | | | | | | | P *vs.* F | 1.21 | $F_{(3, 36)}$ = 1.00; n.s. | — | — |
| | | | | | | | P *vs.* M | 2.41*** | *Pans* | | |
| | | | | | | *Spring* | | | $F_{(3, 36)}$ = = 5.85** | Su *vs.* Sp | 0.08 |
| | | | | | | $F_{(2, 27)}$ = 13.21*** | P vs. F | 0.71 | | Su *vs.* Wi | -0.42 |
| | | | | | | | M vs. F | -1.01* | | Au *vs.* Sp | -0.62 |
| | | | | | | | P vs.M | 1.72*** | | Wi *vs.* Sp | 0.51 |
| | | | | | | | | | | Su *vs.* Au | 0.70 |
| | | | | | | | | | | Au *vs.* Wi | -1.12** |
| Juveniles | Summer | 33±7 | 9±2 | 11±3 | 17±3 | $F_{(2, 106)}$ = 3.17* | M *vs* . F | -2.47* | $F_{(3, 106)}$ = 25.10*** | Au *vs.* Sp | -7.86*** |
| | Autumn | 21±6 | 25±14 | 7±1 | 18±5 | | P *vs.* M | 1.63 | | Su *vs.* Sp | -7.11*** |
| | Winter | 33±8 | 42±17 | 29±8 | 35±7 | | P *vs.* F | -0.85 | | Wi *vs.* Sp | -5.02*** |
| | Spring | 134±29 | 89±31 | 213±30 | 145±19 | | | | | Au *vs.* Wi | -2.77* |
| | | | | | | | | | | Su *vs.* Wi | 2.02 |
| | *Habitats pooled* | 42±10 | 39±9 | 54±13 | | | | | | Su *vs.* Au | 0.45 |

Φ As there was a significant interaction between Habitat and Season when testing density of adult snails, we report effect sizes separately for this variable.
∗ $P$ < 0.05; ∗∗ $P$ < 0.01; ∗∗∗ $P$ < 0.001. We applied the Tukey-HSD correction for multiple testing when comparing effect sizes.

**Table S2.** Results of principal components analyses (PCAs) of shell and aperture morphometric of *Heleobia australis* collected from high and low environmental stressful habitats from the Bahía Blanca estuary, Argentina. Shell size estimated as the volume of a double cone ($mm^3$), shell shape as the SL to SW ratio, aperture size as the area of an ellipse ($mm^2$), and aperture shape as the ratios between aperture length and width (AL to AW ratio).

| Snail groups | Parameters | PC1 | PC2 | PC3 | PC4 |
|---|---|---|---|---|---|
| Infected *vs*. Uninfected | Eigenvalue | 1.404 | 1.086 | 0.837 | 0.385 |
| | Explained variance (%) | 49.3 | 29.5 | 17.5 | 3.7 |
| | Component loadings | | | | |
| | Shell size | -0.662 | 0.228 | -0.030 | 0.714 |
| | Shell shape | -0.245 | -0.683 | -0.687 | -0.039 |
| | Aperture size | -0.639 | 0.322 | -0.052 | -0.697 |
| | Aperture shape | 0.307 | 0.615 | -0.724 | 0.057 |
| Uninfected adults | Eigenvalue | 1.477 | 0.992 | 0.829 | 0.385 |
| | Explained variance (%) | 54.6 | 24.6 | 17.10 | 3.7 |
| | Component loadings | | | | |
| | Shell size | -0.610 | 0.334 | 0.069 | -0.715 |
| | Shell shape | -0.340 | -0.690 | 0.638 | 0.029 |
| | Aperture size | -0.595 | 0.392 | 0.075 | 0.698 |
| | Aperture shape | 0.398 | 0.509 | 0.763 | -0.028 |
| Infected adults | Eigenvalue | 1.366 | 1.069 | 0.920 | 0.382 |
| | Explained variance (%) | 46.7 | 28.6 | 21.1 | 3.6 |
| | Component loadings | | | | |
| | Shell size | 0.703 | -0.049 | -0.037 | -0.708 |
| | Shell shape | 0.020 | 0.720 | -0.694 | 0.005 |
| | Aperture size | 0.700 | -0.091 | -0.069 | 0.705 |
| | Aperture shape | -0.122 | -0.686 | -0.716 | -0.037 |
| Unifected juveniles | Eigenvalue | 1.595 | 0.923 | 0.742 | 0.719 |
| | Explained variance (%) | 63.6 | 21.3 | 13.8 | 1.3 |
| | Component loadings | | | | |
| | Shell size | -0.594 | 0.111 | -0.344 | 0.719 |
| | Shell shape | -0.458 | 0.324 | 0.827 | -0.033 |
| | Aperture size | -0.586 | 0.122 | -0.400 | -0.694 |
| | Aperture shape | -0.307 | -0.931 | 0.194 | -0.016 |

**Table S3. Uninfected adult snails.** Detailed results of linear models on shell characters measured on uninfected adult individuals of the mud snail *Heleobia australis* collected from the intertidal area of the Bahía Blanca estuary, Argentina. In all analyses, we ran models with a Gaussian error distribution and the number of observations was 2,633. Significant values of each effect are indicated in bold.

| Variables | | | | | | Fixed effects | Random effect |
|---|---|---|---|---|---|---|---|
| | | | Habitat | Sex | Habitat : Sex | Shell Volume | Season |
| Shell | Size | | | | $X^2_2$ = 8.42<br>*p*-value **= 0.015** | | variance = 0.270<br>*p*-value **<0.001** |
| | | Male (n=1,254) | $X^2_2$ = 346.22<br>*p*-value **<0.001** | | | | variance = 0.237<br>*p*-value **<0.001** |
| | | Female (n=1,379) | $X^2_2$ = 249.33<br>*p*-value **<0.001** | | | | variance = 0.314<br>*p*-value **<0.001** |
| | | Flats (n=784) | | $X^2_1$ = 0.20<br>*p*-value =0.654 | | | variance = 0.962<br>*p*-value **<0.001** |
| | | Marshes (n=511) | | $X^2_1$ = 1.57<br>*p*-value =0.211 | | | variance = 0.039<br>*p*-value **= 0.008** |
| | | Pans (n=1,338) | | $X^2_1$ = 24.30<br>*p*-value **<0.001** | | | variance = 0.573<br>*p*-value **<0.001** |
| | Shape | | | | $X^2_2$ = 8.20<br>*p*-value **= 0.017** | | variance = 7.6e-05<br>*p*-value **= 0.023** |
| | | Male | $X^2_2$ = 158.77<br>*p*-value **<0.001** | | | | variance = 7.3e-05<br>*p*-value = 0.116 |
| | | Female | $X^2_2$ = 84.03<br>*p*-value **<0.001** | | | | variance = 6.1e-0.5<br>*p*-value **<0.001** |
| | | Flats | | $X^2_1$ = 1.07<br>*p*-value =0.300 | | | variance =1.1e-0.3<br>*p*-value **<0.001** |
| | | Marshes | | $X^2_1$ = 4.15<br>*p*-value **= 0.042** | | | variance = 1.9e-04<br>*p*-value = 0.074 |
| | | Pans | | $X^2_1$ = 37.51<br>*p*-value **<0.001** | | | variance = 2.9e-04<br>*p*-value **= 0.006** |
| | (shell volume as covariate) | | $X^2_2$ = 101.02<br>*p*-value **<0.001** | $X^2_1$ = 29.02<br>*p*-value **<0.001** | $X^2_2$ = 5.69<br>*p*-value = 0.058 | $X^2_1$ = 82.39<br>*p*-value **<0.001** | variance = 2.1e-04<br>*p*-value **= 0.006** |
| Aperture | Size | | $X^2_2$ = 326.62<br>*p*-value **<0.001** | $X^2_1$ = 8.54<br>*p*-value **= 0.003** | $X^2_2$ = 3.05<br>*p*-value = 0.217 | | variance = 0.016<br>*p*-value **<0.001** |
| | Shape | | $X^2_2$ = 535.73<br>*p*-value **<0.001** | $X^2_1$ = 0.04<br>*p*-value = 0.840 | $X^2_2$ = 0.65<br>*p*-value = 0.724 | | variance = 1.19e-03<br>*p*-value **<0.001** |
| | (shell volume as covariate) | | $X^2_2$ = 287.66<br>*p*-value **<0.001** | $X^2_1$ = 0.70<br>*p*-value = 0.401 | $X^2_2$ = 2.20<br>*p*-value = 0.333 | $X^2_1$ = 134.7<br>*p*-value **<0.001** | variance = 1.21e-03<br>*p*-value **<0.001** |

**Table S4. Infected adult snails.** Summary of means (±SE) and statistical significance of habitat and sex effect on shell and aperture morphometric of infected adult individuals of the mud snail *Heleobia australis* from the intertidal area of the Bahía Blanca estuary, Argentina. Variables are the same as indicated in table 3. Number of observations are indicated between parentheses, which are only shown for shell size but are the same for other traits measured.

| Traits measured | Habitat type | Sex | | *Sex pooled* | Habitat effect | Habitat comparison | Effect size | Sex Effect | Sex comparison | Effect size |
|---|---|---|---|---|---|---|---|---|---|---|
| | | Male (♂) | Female (♀) | | | | | | | |
| Shell size | Flats (F) | 6.33±0.16 (78) | 6.02±0.15 (98) | 6.16±0.11 (176) | $X_2^2$ = 37.40*** | P *vs.* F | 5.96*** | $X_1^2$ = 11.38*** | ♂ *vs.* ♀ | 3.39*** |
| | Marshes (M) | 6.77±0.27 (40) | 6.56±0.97 (73) | 6.62±0.75 (113) | | P *vs.* M | 2.74* | | | |
| | Pans (P) | 7.01±0.14 (432) | 6.76±0.20 (573) | 6.87±0.16 (1005) | | M *vs* . F | 1.86 | | | |
| | *Habitats pooled* | 6.88±0.11 (550) | 6.56±0.17 (744) | | | | | | | |
| Shell shape | Flats | 2.32±0.02 | 2.29±0.02 | 2.31±0.01 | $X_2^2$ = 54.41*** | P *vs.* F | 6.68*** | $X_1^2$ = 0.33; n.s. | ♂ *vs.* ♀ | 0.55 |
| | Marshes | 2.34±0.02 | 2.32±0.02 | 2.32±0.02 | | P *vs.* M | 4.44*** | | | |
| | Pans | 2.40±0.14 | 2.39±0.02 | 2.40±0.01 | | M *vs* . F | 0.92 | | | |
| | *Habitats pooled* | 2.38±0.01 | 2.37±0.02 | | | | | | | |
| Aperture size | Flats | 1.72±0.04 | 1.65±0.04 | 1.68±0.02 | $X_2^2$ = 18.14*** | P *vs.* F | 3.89*** | $X_1^2$ = 8.89** | ♂ *vs.* ♀ | 2.99** |
| | Marshes | 1.76±0.06 | 1.63±0.06 | 1.68±0.03 | | P *vs.* M | 2.48* | | | |
| | Pans | 1.80±0.04 | 1.76±0.06 | 1.77±0.05 | | M *vs* . F | 0.62 | | | |
| | *Habitats pooled* | 1.78±0.03 | 1.72±0.05 | | | | | | | |
| Aperture shape | Flats | 1.69±0.03 | 1.66±0.03 | 1.67±0.03 | $X_2^2$ = 157.24*** | P *vs.* F | -11.54*** | $X^2_1$ = 5e-04; n.s. | ♂ *vs.* ♀ | -0.84 |
| | Marshes | 1.65±0.04 | 1.66±0.03 | 1.66±0.03 | | P *vs.* M | -7.86*** | | | |
| | Pans | 1.56±0.02 | 1.56±0.02 | 1.56±0.02 | | M *vs* . F | -1.42 | | | |
| | *Habitats pooled* | 1.58±0.03 | 1.58±0.03 | | | | | | | |

Estimates for shell and aperture shape are uncorrected as they do not change when removing the covariate from the model.
$*$ $P < 0.05$; $**$ $P < 0.01$; $***$ $P < 0.001$. We applied the Holm-Bonferroni correction for multiple testing when comparing effect sizes.

**Table S5. Infected adult snails**. Detailed results of linear models on shell characters measured on infected adult individuals of the mud snail *Heleobia australis* collected from the intertidal area of the Bahía Blanca estuary, Argentina. In all analyses, we ran models with a Gaussian error distribution and the number of observations was 1,294. Significant values of each effect are indicated in bold.

| Variables | | Fixed effects | | | | Random effect |
|---|---|---|---|---|---|---|
| | | Habitat | Sex | Habitat : Sex | Shell Volume | Season |
| Shell | Size | $X^2_2 = 37.40$ | $X^2_1 = 11.38$ | $X^2_2 = 0.43$ | | variance = 0.09 |
| | | *p* -value **<0.001** | *p* -value **<0.001** | *p* -value = 0.807 | | *p* -value **<0.001** |
| | Shape | $X^2_2 = 54.41$ | $X^2_1 = 0.33$ | $X^2_2 = 2.64$ | | variance = 4.26e-04 |
| | | *p* -value **<0.001** | *p* -value = 0.560 | *p* -value = 0.267 | | *p* -value **<0.001** |
| | (shell volume as covariate) | $X^2_2 = 58.78$ | $X^2_1 = 0.38$ | $X^2_2 = 2.66$ | $X^2_1 = 0.26$ | variance = 4.13e-04 |
| | | *p* -value **<0.001** | *p* -value = 0.536 | *p* -value = 0.265 | *p* -value = 0.689 | *p* -value **<0.001** |
| Aperture | Size | $X^2_2 = 18.14$ | $X^2_1 = 8.89$ | $X^2_2 = 1.22$ | | variance = 7.0e-04 |
| | | *p* -value **<0.001** | *p* -value **= 0.003** | *p* -value = 0.542 | | *p* -value **<0.001** |
| | Shape | $X^2_2 = 157.24$ | $X^2_1 = 0.04$ | $X^2_2 = 0.97$ | | variance = 1.63e-03 |
| | | *p* -value **<0.001** | *p* -value = 0.850 | *p* -value = 0.616 | | *p* -value **<0.001** |
| | (shell volume as covariate) | $X^2_2 = 145.11$ | $X^2_1 = 5e-04$ | $X^2_2 = 0.92$ | $X^2_1 = 4.98$ | variance = 1.68e-03 |
| | | *p* -value **<0.001** | *p* -value = 0.982 | *p* -value = 0.631 | *p* -value **= 0.026** | *p* -value **<0.001** |

**Table S6. Uninfected juvenile snails.** Summary of means (±SE) and statistical significance of sex effect on shell and aperture morphometric of uninfected juveniles of the mud snail *Heleobia australis* from the intertidal area Bahía Blanca estuary, Argentina. Variables are the same as indicated in table 3. Number of observations are indicated between parentheses, which are only shown for shell size but are the same for other traits measured.

| Traits measured | Habitat type | Sex | | *Sex pooled* | Habitat effect | Habitat comparison | Effect size | Sex Effect | Sex comparison | Effect size |
|---|---|---|---|---|---|---|---|---|---|---|
| | | Male (♂) | Female (♀) | | | | | | | |
| Shell size Φ | Flats (F) | 3.25±0.08 (81) | 1.76±0.21 (964) | 1.85±0.23 (1045) | *Males* | P *vs.* F | 4.61*** | *Flats* | | |
| | Marshes (M) | 3.50±0.21 (61) | 1.84±0.19 (674) | 2.01±0.22 (735) | $X_2^2$ = 22.35*** | P *vs.* M | 3.42** | $X_1^2$ = 193.75*** | ♂ *vs.* ♀ | 13.92*** |
| | Pans (P) | 3.95±0.11 (70) | 2.69±0.20 (189) | 2.91±0.24 (259) | | M *vs* . F | 1.07 | *Marshes* | | |
| | *Habitats pooled* | 3.52±0.13 (212) | 1.87±0.15 (1489) | | *Females* | P *vs.* F | 13.71*** | $X_1^2$ = 181.94*** | ♂ *vs.* ♀ | 13.49*** |
| | | | | | $X_2^2$ = 207.51*** | P *vs.* M | 13.56*** | *Pans* | | |
| | | | | | | M *vs* . F | 0.49 | $X_1^2$ = 43.69*** | ♂ *vs.* ♀ | 4.93*** |
| Shell shape | Flats | 2.05±0.02 | 1.95±0.02 | 1.95±0.02 | $X_2^2$ = 264.74*** | P *vs.* F | 15.85*** | $X_1^2$ = 126.65*** | ♂ *vs.* ♀ | 11.40*** |
| | Marshes | 2.05±0.01 | 1.96±0.01 | 1.96±0.01 | | P *vs.* M | 15.43*** | | | |
| | Pans | 2.11±0.01 | 2.03±0.02 | 2.06±0.02 | | M *vs* . F | 1.03 | | | |
| | *Habitats pooled* | 2.07±0.01 | 1.96±0.01 | | | | | | | |
| Aperture size Φ | Flats | 1.23±0.03 | 0.85±0.07 | 0.87±0.07 | *Males* | P *vs.* F | 2.90* | *Flats* | | |
| | Marshes | 1.21±0.03 | 0.84±0.03 | 0.88±0.04 | $X_2^2$ = 9.60** | P *vs.* M | 2.52* | $X_1^2$ = 130.58*** | ♂ *vs.* ♀ | 11.43*** |
| | Pans | 1.32±0.05 | 1.09±0.06 | 1.13±0.07 | | M *vs* . F | 0.51 | *Marshes* | | |
| | *Habitats pooled* | 1.26±0.04 | 0.87±0.04 | | *Females* | P *vs.* F | 12.44*** | $X_1^2$ = 100.11*** | ♂ *vs.* ♀ | 10.01*** |
| | | | | | $X_2^2$ = 165.59*** | P *vs.* M | 11.88*** | *Pans* | | |
| | | | | | | M *vs* . F | 1.50 | $X_1^2$ = 124.81*** | ♂ *vs.* ♀ | 4.98*** |
| Aperture shape † | Flats | 1.69±0.02 | 1.67±0.02 | 1.67±0.02 | $X_2^2$ = 52.42*** | P *vs.* F | -2.04 | $X_1^2$ = 3.94; n.s. | ♂ *vs.* ♀ | -1.99 |
| | Marshes | 1.64±0.02 | 1.63±0.01 | 1.63±0.01 | | P *vs.* M | -1.67 | | | |
| | Pans | 1.61±0.01 | 1.61±0.02 | 1.61±0.02 | | M *vs* . F | -0.17 | | | |
| | *Habitats pooled* | 1.65±0.02 | 1.65±0.02 | | | | | | | |

Φ As there was a significant interaction between Habitat and Sex when testing shell and aperture size, we report effect sizes separately for these traits.
† We report estimates of shell shape corrected for shell size, as the pattern observed was not the same when removing the covariate from the model.
∗ $P < 0.05$; ∗∗ $P < 0.01$; ∗∗∗ $P < 0.001$. We applied the Holm-Bonferroni correction for multiple testing when comparing effect sizes.

**Table S7. Uninfected juvenile snails.** Detailed results of linear models on shell characters measured on uninfected juvenile individuals of the mud snail *Heleobia australis* collected from the intertidal area of the Bahía Blanca estuary, Argentina. In all analyses, we ran models with a Gaussian error distribution and the number of observations was 2,039. Significant values of each effect are indicated in bold.

| Variables | | | Fixed effects | | | | Random effect |
|---|---|---|---|---|---|---|---|
| | | | Habitat | Sex | Habitat : Sex | Shell Volume | Season |
| Shell | Size | | | | $X^2_2 = 8.22$<br>*p*-value = **0.016** | | variance = 0.06<br>*p*-value **<0.001** |
| | | Male (n=212) | $X^2_2 = 22.35$<br>*p*-value **<0.001** | | | | variance = 0.015<br>*p*-value = 0.139 |
| | | Female (n=1,827) | $X^2_2 = 207.51$<br>*p*-value **<0.001** | | | | variance = 0.064<br>*p*-value **<0.001** |
| | | Flats (n=1,045) | | $X^2_1 = 193.75$<br>*p*-value **<0.001** | | | variance = 0.159<br>*p*-value **<0.001** |
| | | Marshes (n=735) | | $X^2_1 = 181.94$<br>*p*-value **<0.001** | | | variance = 0.141<br>*p*-value **<0.001** |
| | | Pans (n=259) | | $X^2_1 = 43.69$<br>*p*-value **<0.001** | | | variance = 0.105<br>*p*-value **<0.001** |
| | Shape | | $X^2_2 = 157.95$<br>*p*-value **<0.001** | $X^2_1 = 135.34$<br>*p*-value **<0.001** | $X^2_2 = 1.48$<br>*p*-value = 0.477 | | variance = 5.3e-04<br>*p*-value **<0.001** |
| | (shell volume as covariate) | | $X^2_2 = 38.87$<br>*p*-value **<0.001** | $X^2_1 = 5.12$<br>*p*-value **= 0.024** | $X^2_2 = 0.01$<br>*p*-value = 0.994 | $X^2_1 = 483.4$<br>*p*-value **<0.001** | variance = 6.2e-04<br>*p*-value **<0.001** |
| Aperture | Size | | | | $X^2_2 = 9.93$<br>*p*-value = **0.007** | | variance = 4.3e-03<br>*p*-value **<0.001** |
| | | Male | $X^2_2 = 9.60$<br>*p*-value **= 0.008** | | | | variance = 2.4e-03<br>*p*-value **= 0.017** |
| | | Female | $X^2_2 = 165.59$<br>*p*-value **<0.001** | | | | variance =5.0e-03<br>*p*-value **<0.001** |
| | | Flats | | $X^2_1 = 130.58$<br>*p*-value **<0.001** | | | variance = 1.7e-03<br>*p*-value **<0.001** |
| | | Marshes | | $X^2_1 = 100.11$<br>*p*-value **<0.001** | | | variance = 3.0e-03<br>*p*-value **<0.001** |
| | | Pans | | $X^2_1 = 24.81$<br>*p*-value **<0.001** | | | variance = 1.1e-02<br>*p*-value **<0.001** |
| | Shape | | $X^2_2 = 20.03$<br>*p*-value **<0.001** | $X^2_1 = 62.89$<br>*p*-value **<0.001** | $X^2_2 = 1.89$<br>*p*-value = 0.389 | | variance = 1.4e-03<br>*p*-value **<0.001** |
| | (shell volume as covariate) | | $X^2_2 = 52.50$<br>*p*-value **<0.001** | $X^2_1 = 2.06$<br>*p*-value = 0.151 | $X^2_2 = 1.70$<br>*p*-value = 0.426 | $X^2_1 = 206.1$<br>*p*-value **<0.001** | variance = 8.6e-04<br>*p*-value **<0.001** |

**Table S8. Infected vs. uninfected adult snails.** Detailed results of linear models on shell characters measured on infected and uninfected adult individuals of the mud snail *Heleobia australis* collected from the intertidal area of the Bahía Blanca estuary, Argentina. In all analyses, we ran models with a Gaussian error distribution and the number of observations was 2,343. Significant values of each effect are indicated in bold. Note that only individuals from pans were considered, which allowed for removing the effect of habitat conditions on shell morphometrics.

| Variables | | | Fixed effects | | | | Random effect |
|---|---|---|---|---|---|---|---|
| | | | Status | Sex | Status : Sex | Shell Volume | Season |
| Shell | Size | | $X^2_1$ = 149.69<br>*p*-value **<0.001** | $X^2_1$ = 31.68<br>*p*-value **<0.001** | $X^2_1$ = 0.70<br>*p*-value = 0.402 | | variance = 0.312<br>*p*-value **<0.001** |
| | Shape | | | | $X^2_1$ = 15.86<br>*p*-value **<0.001** | | variance = 1.1e-04<br>*p*-value **= 0.012** |
| | | Male (n=1,032) | $X^2_1$ = 11.82<br>*p*-value **<0.001** | | | | variance = 2.0e-04<br>*p*-value **= 0.048** |
| | | Female (n=1,311) | $X^2_1$ = 99.50<br>*p*-value **<0.001** | | | | variance = 4.9e-05<br>*p*-value = 0.200 |
| | | Unifected (n=1,338) | | $X^2_1$ = 37.51<br>*p*-value **<0.001** | | | variance = 2.9e-04<br>*p*-value **= 0.006** |
| | | Infected (n=1,005) | | $X^2_1$ = 0.09<br>*p*-value = 0.766 | | | variance = 5.7e-04<br>*p*-value **<0.001** |
| | (shell volume as covariate) | | | | $X^2_1$ = 15.95<br>*p*-value **<0.001** | $X^2_1$ = 11e03<br>*p*-value **<0.001** | variance = 1.2e-04<br>*p*-value **= 0.019** |
| | | Male | $X^2_1$ = 10.21<br>*p*-value **= 0.001** | | | $X^2_1$ = 0.23<br>*p*-value = 0.630 | variance = 2.0e-03<br>*p*-value = 0.052 |
| | | Female | $X^2_1$ = 87.47<br>*p*-value **<0.001** | | | $X^2_1$ = 1.84<br>*p*-value = 0.175 | variance = 2.6e-05<br>*p*-value = 0.317 |
| | | Unifected | | $X^2_1$ = 35.60<br>*p*-value **<0.001** | | $X^2_1$ = 0.60<br>*p*-value = 0.438 | variance = 2.6e-04<br>*p*-value **= 0.012** |
| | | Infected | | $X^2_1$ = 0.001<br>*p*-value = 0.975 | | $X^2_1$ = 9.35<br>*p*-value **= 0.002** | variance = 4.4e-04<br>*p*-value **<0.001** |
| Aperture | Size | | $X^2_1$ = 408.65<br>*p*-value **<0.001** | $X^2_1$ = 14.95<br>*p*-value **<0.001** | $X^2_1$ = 2e-04<br>*p*-value = 0.990 | | variance = 0.020<br>*p*-value **<0.001** |
| | Shape | | $X^2_1$ = 60.57<br>*p*-value **<0.001** | $X^2_1$ = 1.3e-3<br>*p*-value = 0.971 | $X^2_1$ = 0.33<br>*p*-value = 0.564 | | variance = 1.5e-03<br>*p*-value **<0.001** |
| | (shell volume as covariate) | | $X^2_1$ = 91.79<br>*p*-value **<0.001** | $X^2_1$ = 0.92<br>*p*-value = 0.337 | $X^2_1$ = 0.52<br>*p*-value = 0.470 | $X^2_1$ = 62.85<br>*p*-value **<0.001** | variance = 1.6e-03<br>*p*-value **<0.001** |

**Table S9. Shell and body weight.** Detailed results of linear models on shell and body weight measured on individuals of the mud snail *Heleobia australis* collected from the intertidal area of the Bahía Blanca estuary, Argentina. In both analyses, we ran models with a Gaussian error distribution and the number of observations was 814. Significant values of each effect are indicated in bold. In this study, shell weight was used as a proxy of shell thickness.

| Variables | Fixed effects | | Random effect |
|---|---|---|---|
| | Habitat | Shell volume | Season |
| Shell weight | $X^2_2$ = 48.93 | $X^2_1$ = 3238.57 | variance = 0.06 |
| | *p*-value **<0.001** | *p*-value **<0.001** | *p*-value **<0.001** |
| Ln(Body weight) | $X^2_2$ = 19.67 | $X^2_1$ = 99.74 | variance = 1.90E-0.3 |
| | *p*-value **<0.001** | *p*-value **<0.001** | *p*-value=0.083 |

### SUPPLEMENTARY FIGURES

**Figure S1.** Length-frequency distributions and polymodal decomposition of individuals of each subpopulation of the snail *Heleobia australis* collected in 2012 from the Bahía Blanca estuary, Argentina. Cohort 1, cohort of individuals from second year age-class (Adults); Cohort 2, cohort of individuals from first year age-class (Juveniles); SL, mean shell-length (± SD); n, number of individiduals in each cohort.

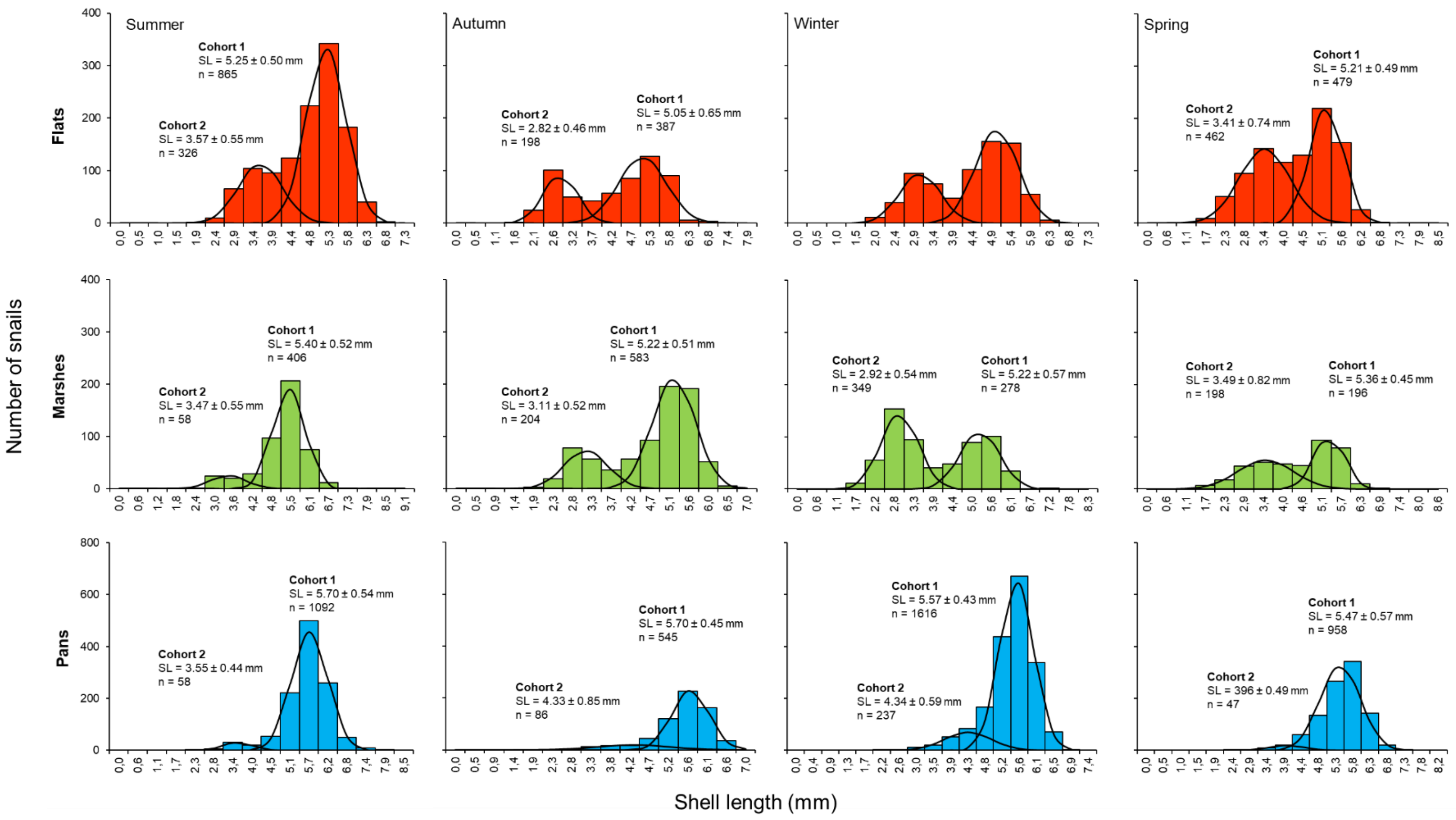

**Figure S2.** Variation in juvenile and adult snail density of *Heleobia australis* across habitats and seasons in the intertidal area of the Bahía Blanca estuary, Argentina. Bars represent ± 1 SE.

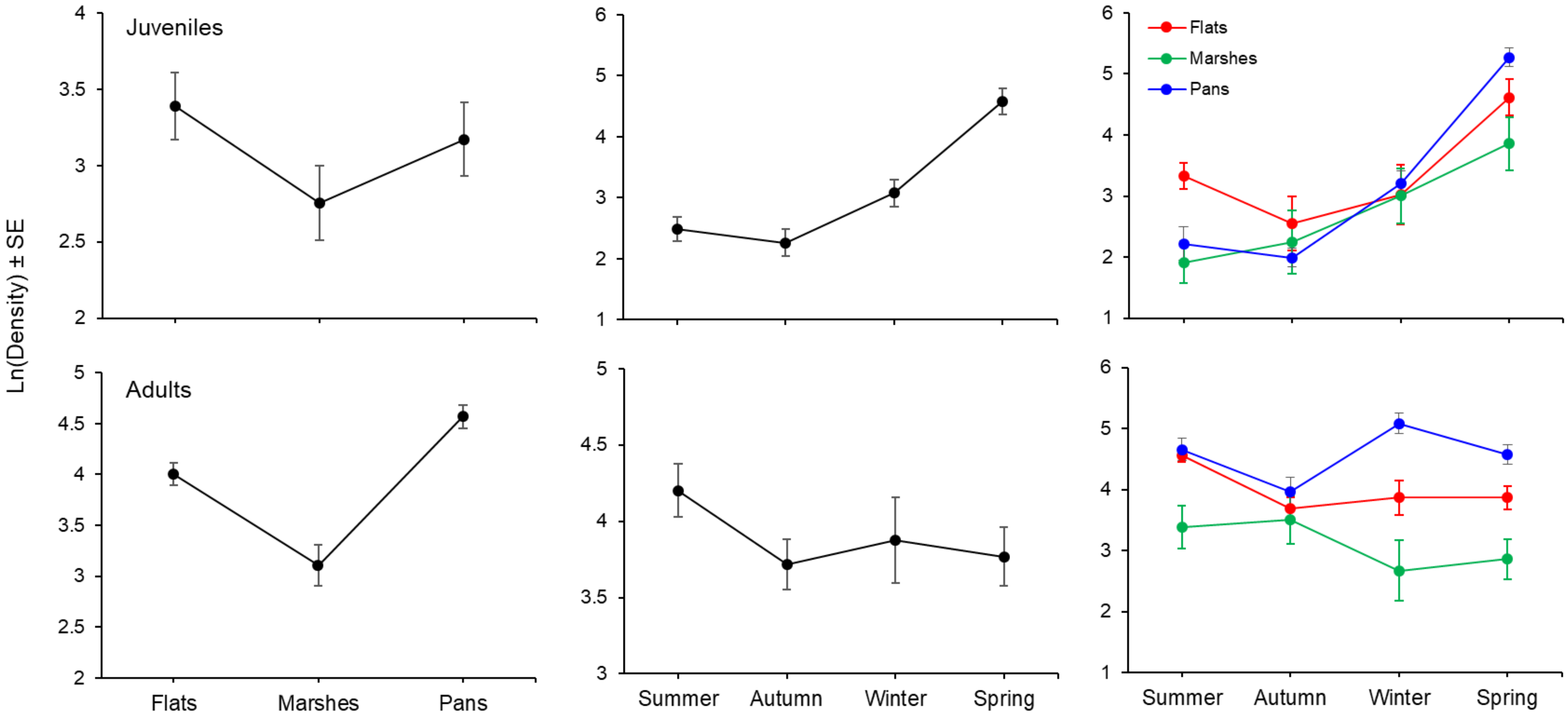

**Figure S3.** Biplots of principal component factor scores, factor loadings, and normal confidence ellipses (95%) of individual measurements of shell characteristics (shell and aperture size and shape) of snails from habitats with high (flats and marshes) or low (pans) environmental stress conditions: uninfected adults (cat. *i*), infected adults (cat. *ii*), uninfected juveniles (cat. *iii*) and with different infection condition (uninfected vs. infected adult snails; cat. *iv*).

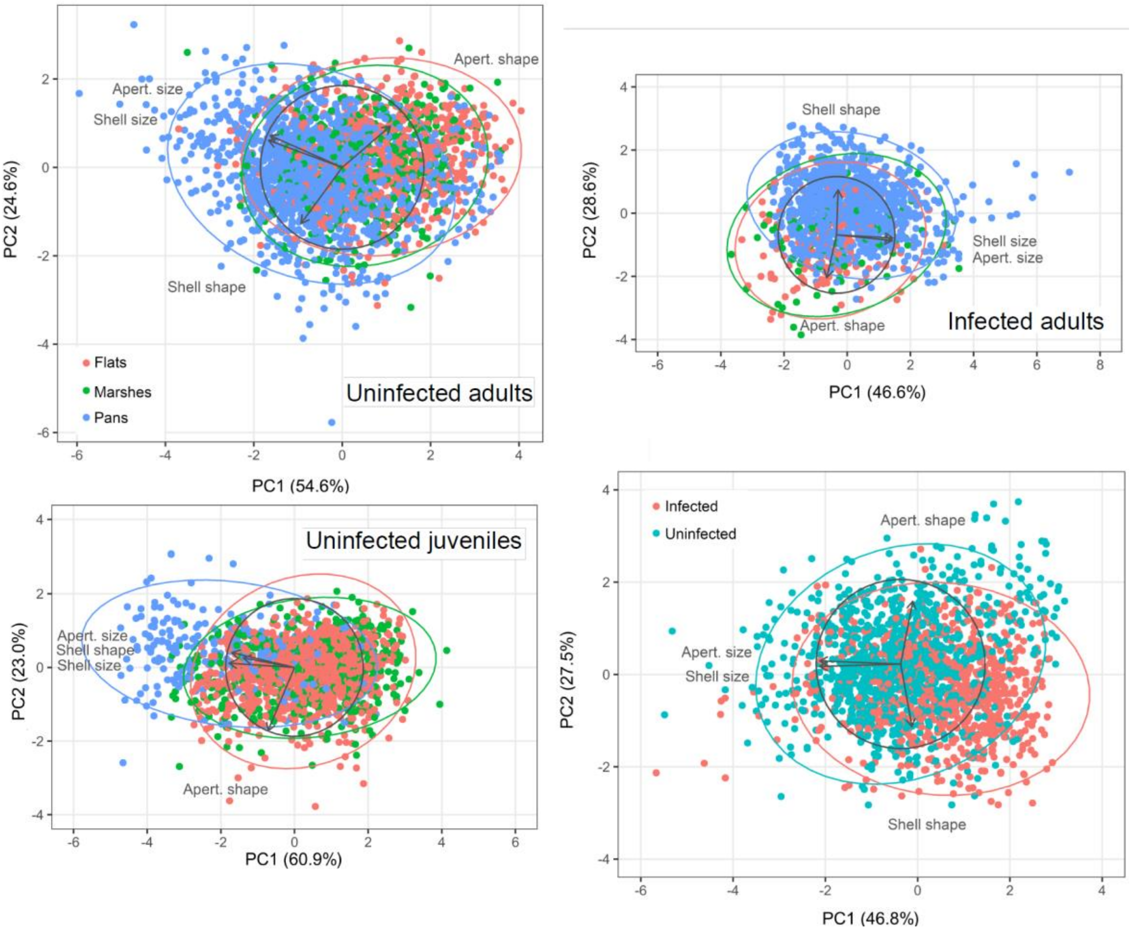

**Figure S4. Infected adult snails (cat. *ii*).** Density plots of habitat effect on shell and aperture morphometrics of uninfected juvenile snails *Heleobia australis* in the Bahía Blanca estuary, Argentina. Blue and red dots indicate mean values of each variable and sex in each habitat (flats, marshes, and pans). Bars represent ± 1 SE.

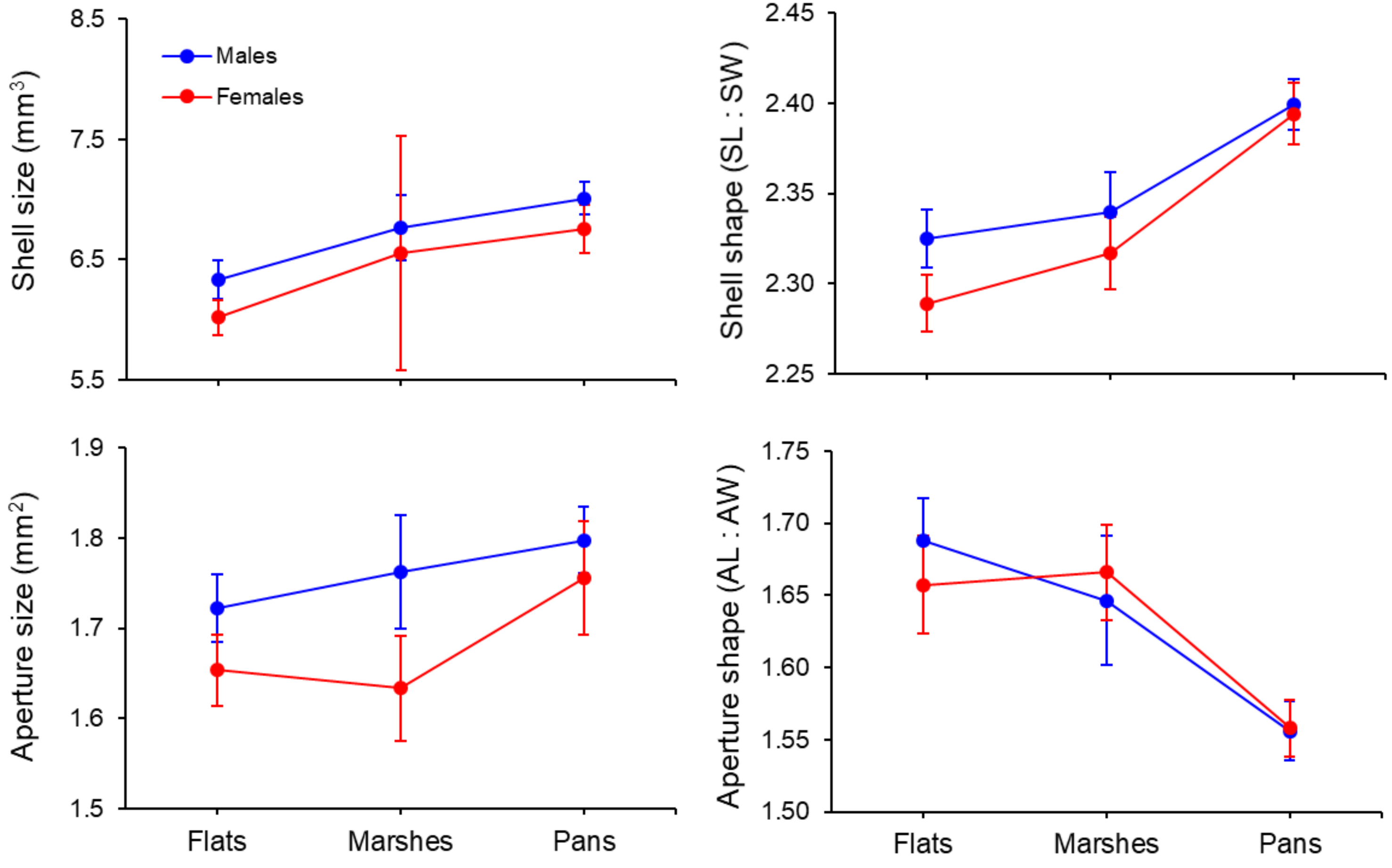

**Figure S5. Uninfected juveniles snails (cat. *iii*).** Habitat effect on shell and aperture morphometrics of uninfected juvenile snails *Heleobia australis* in the Bahía Blanca estuary, Argentina. Mean values of aperture shape were statistically corrected for shell size. Blue and red dots indicate mean values of each variable and sex in each habitat (flats, marshes, and pans). Bars represent ± 1 SE.

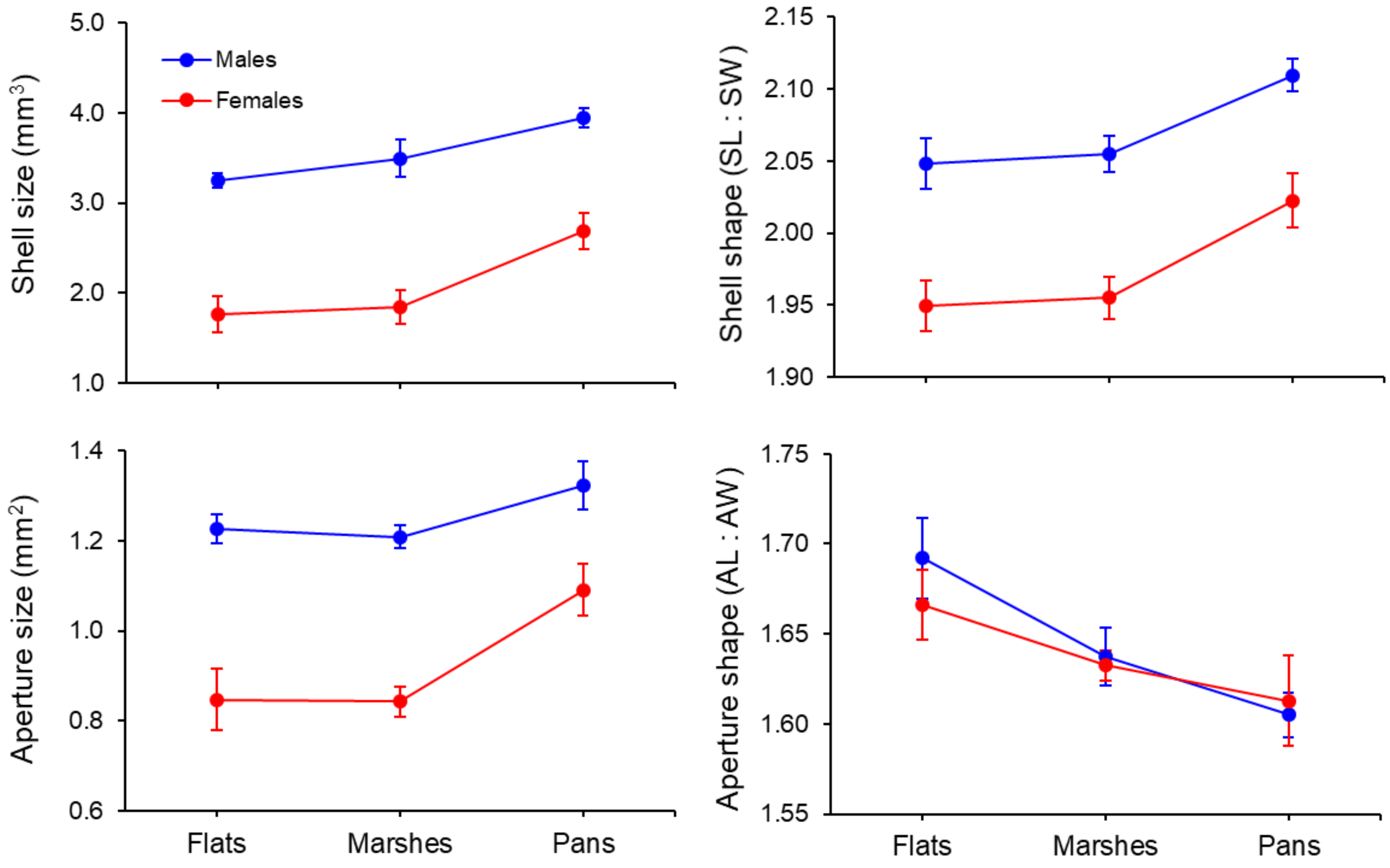

**Figure S6.** Shell weight (used as a proxy of shell thickness) and body mass variation of the intertidal mud snail *Heleobia australis* across habitats with different environmental stressful conditions in the Bahía Blanca estuary, Argentina. Empty diamonds indicate uncorrected mean values whereas black circles show corrected mean values.

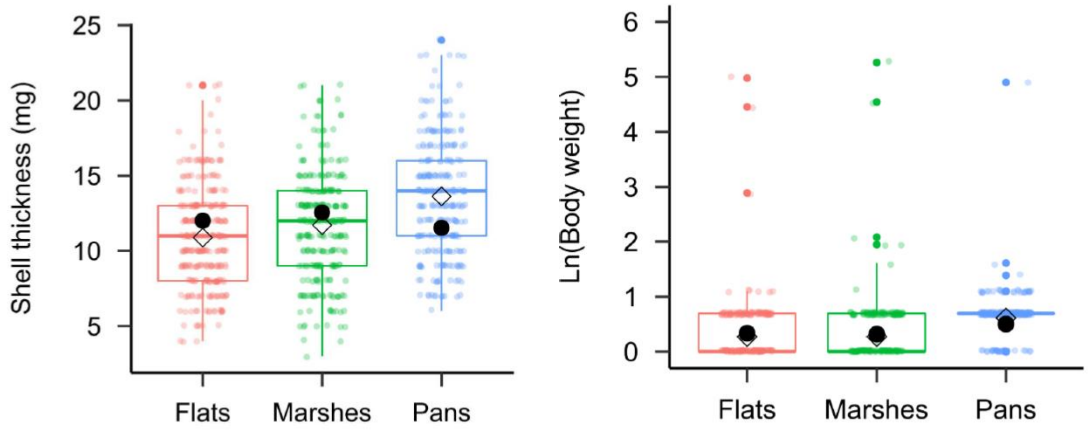

**Figure S7.** Penis morphology of uninfected adult snails *Heleobia australis* collected from three environmentally distinct habitats from the Bahía Blanca estuary, Argentina and kept in standard laboratory conditions for 21 months since they were juveniles, which ensured that individuals analyzed were all adults. Two different views from the penis morphology for each habitat. Bar indicates 1 mm.

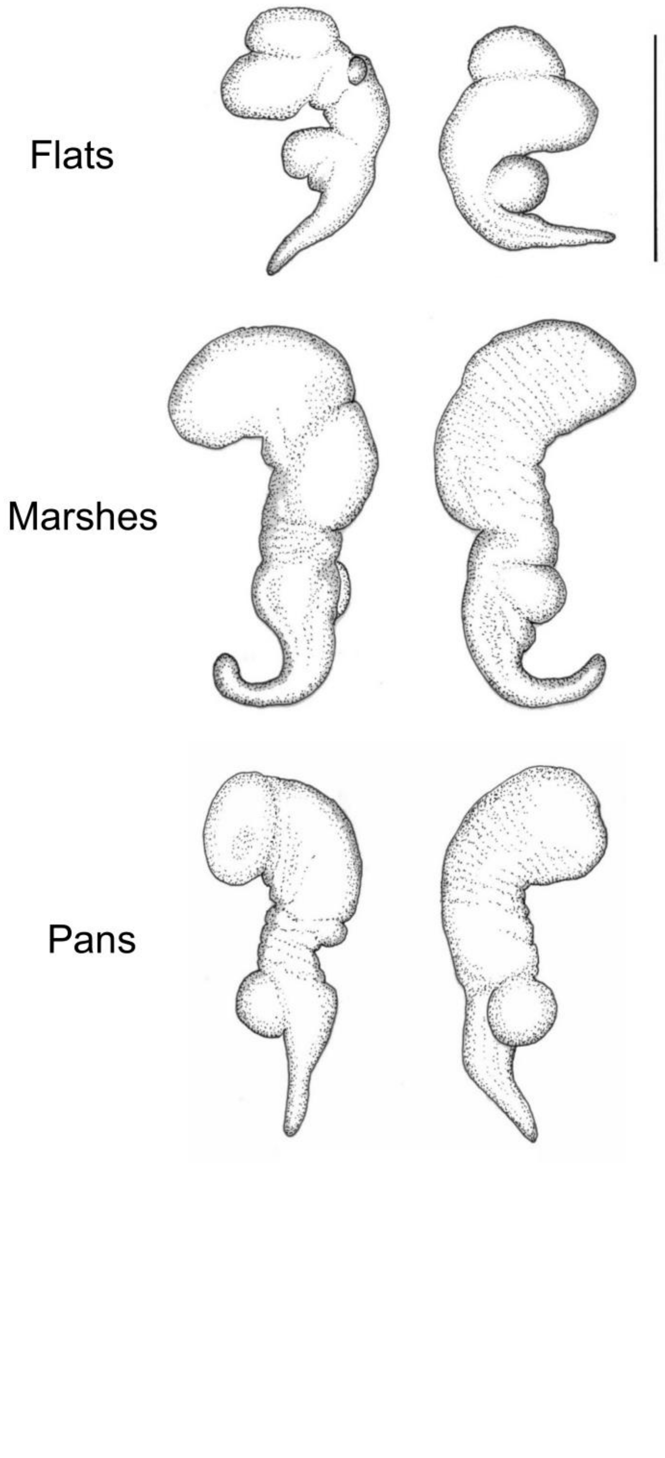

**Figure S8.** Phylogenetic tree of *Heleobia* individuals based on Bayesian inference in Beast2 of the COI gene. All sequences from the current study, as well as the one retrieved from GenBank (yellow), are included in this tree. Numbers indicate posterior probability and sequence coloration represents habitats: flats (pink), marshes (green), and pans (blue).

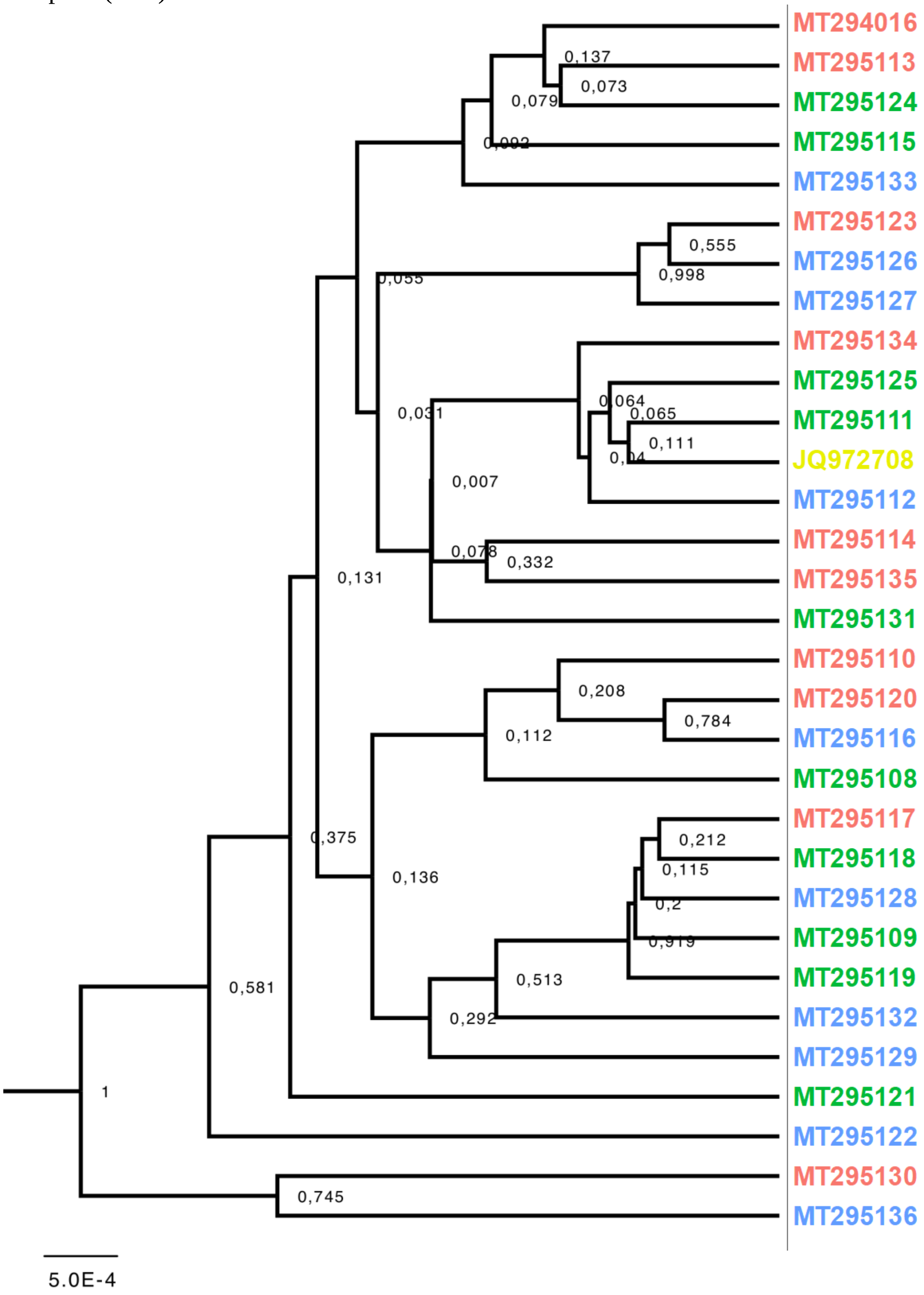

**Figure S9.** Haplotype network of *Heleobia* individuals based on COI gene. Circle sizes are proportional to haplotype frequencies and colors represents location and habitats. The number of mutations separating circles are indicated by dashes.

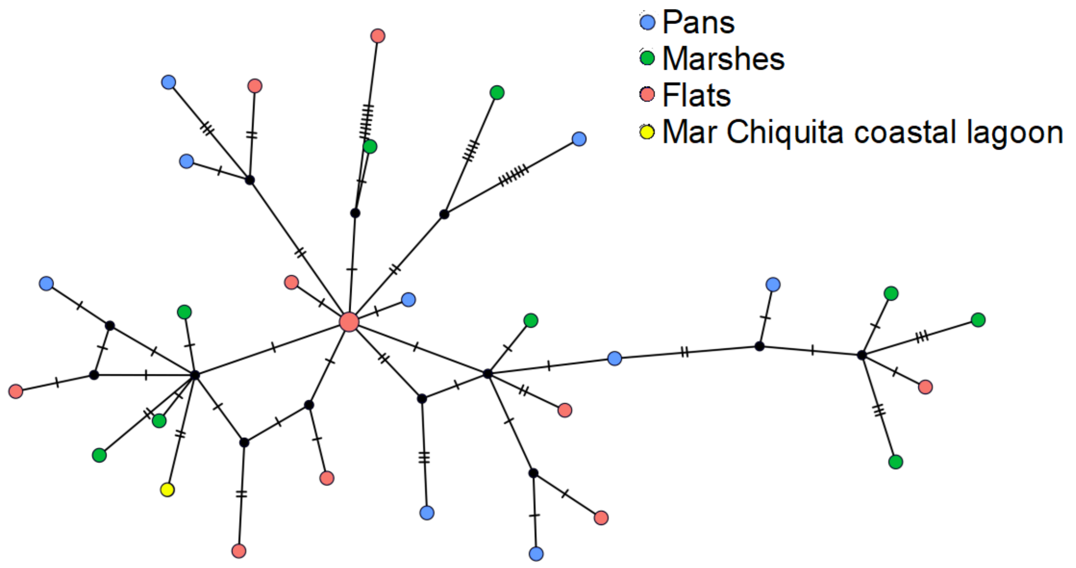
